# Muscarinic receptors regulate auditory and prefrontal cortical communication during auditory processing

**DOI:** 10.1101/285601

**Authors:** Nicholas M. James, Howard J. Gritton, Nancy Kopell, Kamal Sen, Xue Han

**Affiliations:** Boston University, Department of Biomedical Engineering, Boston, MA 02215; Boston University, Department of Mathematics & Statistics, Boston, MA 02215

**Keywords:** Sensory Processing, Auditory Cortex, Prefrontal Cortex, Acetylcholine, Gating

## Abstract

Much of our understanding about how acetylcholine modulates prefrontal cortical (PFC) networks comes from behavioral experiments that examine cortical dynamics during highly attentive states. However, much less is known about how PFC is recruited during passive sensory processing and how acetylcholine may regulate connectivity between cortical areas outside of task performance. To investigate the involvement of PFC and cholinergic neuromodulation in passive auditory processing, we performed simultaneous recordings in the auditory cortex (AC) and PFC in awake head fixed mice presented with a white noise auditory stimulus in the presence or absence of local cholinergic antagonists in AC. We found that a subset of PFC neurons were strongly driven by auditory stimuli even when the stimulus had no associative meaning, suggesting PFC monitors stimuli under passive conditions. We also found that cholinergic signaling in AC shapes the strength of auditory driven response in PFC, by modulating the intra-cortical sensory response through muscarinic interactions in AC. Taken together, these findings provide novel evidence that cholinergic mechanisms have a continuous role in cortical gating through muscarinic receptors during passive processing and expand traditional views of prefrontal cortical function and the contributions of cholinergic modulation in sensory gating.

**Highlights:**

- Prefrontal cortex actively monitors non-associative stimuli under passive conditions
- Acetylcholine facilitates cortical signaling even outside of attentional contexts
- Local scopolamine infusion reduced intracortical signaling and impaired cortical gating
- mAChR have an ongoing role in sound processing

## 1. Introduction

The prefrontal cortex (PFC) in mammals has been associated with various cognitive functions, such as stimulus categorization and integration, decision making, and executive control (Dalley et al., 2004; Miller and Cohen, 2001; Roy et al., 2014). PFC neurons exhibit a high degree of selectivity for conditioned sensory stimuli and can be differentially regulated by contextual associations or task demands (Chang et al., 2010; Euston et al., 2012; Hyman et al., 2012; Maren and Quirk, 2004; Moorman and Aston-Jones, 2015). Interestingly, prefrontal regions are also responsive to conditioned auditory stimuli that had been previously learned, when re-tested under anesthesia, suggesting that learned sensory information can reach PFC in the absence of behavior (Pirch et al., 1985a; Pirch et al., 1985b). While much is known about how PFC is modulated within the context of well-learned behavioral tasks, the intrinsic responsiveness of subregions of the PFC to non-behaviorally relevant sensory stimuli is less well established. In particular, we wanted to know if sensory information is conveyed across multiple prefrontal regions during awake passive states, and is there an underlying mechanism that gates sensory signals into PFC under these conditions?

One candidate for gating sensory input into executive regions is the neurotransmitter acetylcholine, which innervates both sensory and association cortices through projections from the basal forebrain cholinergic system. The cholinergic system is well suited to coordinate neural activity across large brain areas, as it provides diffuse and global innervation of the cortex (Zaborszky et al., 2012). The extensive reach of these projections allow for coincident cholinergic release in disparate regions, which could coordinate networks during stimulus processing, or task engagement by enhancing neural synchrony. Synchrony among distal neural networks has been proposed to facilitate information transfer across interconnected areas (Buschman and Miller, 2007; Fries et al., 2001; Siegel et al., 2012).

Much of our understanding of cholinergic influences on stimulus processing comes from task performing animals where cholinergic agonists generally improve performance or sensory discrimination (Boix-Trelis et al., 2007; Herrero et al., 2008; Liang et al., 2008; Rogers and Kesner, 2004). Studies in which basal forebrain stimulation is paired with concomitant sensory stimuli have demonstrated that activation of cholinergic networks enhances the response reliability of cortical neurons, and increases the correlated temporal activity of individual neurons within the networks they serve (Froemke et al., 2007; Goard and Dan, 2009; Kilgard and Merzenich, 2002; Letzkus et al., 2011; Pinto et al., 2013). While these studies suggest that the cholinergic system has an active role in modulating neural networks during active behavioral states, much less is known about how the cholinergic system contributes to cortical dynamics and cross-regional connectivity in the absence of well-defined behavioral associations.

Previous work has shown that the cholinergic projections densely innervate primary and secondary auditory regions affording them the potential to regulate auditory processing across multiple levels (Chavez and Zaborszky, 2017). Cholinergic modulation of cortical processing occurs via nicotinic (nAChRs) and muscarinic (mAChRs) receptor activation; both of which can facilitate sensory processing in complementary ways. In auditory cortex (AC), nAChRs located on thalamic projection terminals have been shown to augment the strength of presynaptic inputs (Gil et al., 1997; Kawai et al., 2007; Lambe et al., 2003). There is also evidence that post-synaptically located nAChRs can increase excitability by increasing the frequency, magnitude, and duration of the postsynaptic response (Chu et al., 2000; Levy and Aoki, 2002; Roerig et al., 1997). In parallel, acetylcholine has been shown to enhance how the visual cortex responds to stimuli presented within an attended visual receptive field by acting at metabotropic mAChRs (Herrero et al., 2008). mAChRs are ubiquitously expressed across almost all major cortical inhibitory neuron subtypes (Disney and Aoki, 2008), where activation leads to overall enhanced cortical excitability primarily through reducing peri-somatic inhibition of layer V pyramidal neurons (Kruglikov and Rudy, 2008; Nunez et al., 2012). Acetylcholinesterase inhibitors, locally applied, generally mimic the effects of individual agonists on auditory cortex, enhancing frequency specific responses (Ashe et al., 1989). Given their diverse functionality, both nAChRs and mAChRs could coincidently promote cortical representations of sensory inputs through increasing the signal to noise ratio of sensory information (Hasselmo and Sarter, 2011). While several studies have identified a role for cholinergic signaling in modulating cortical representations in auditory cortex during awake or anesthetized conditions (Ashe et al., 1989; Bandrowski et al., 2001; Chu et al., 2000; McKenna et al., 1988; McKenna et al., 1989; Metherate et al., 1990; Metherate et al., 1992; Metherate et al., 2012; Metherate and Weinberger, 1989, 1990), the contributions of cholinergic signaling in conveying sensory representations from auditory to association areas, and the potential contributions of each receptor subtype to this process, has not been explored.

To investigate if PFC monitors sensory stimuli during passive sound processing, and if the cholinergic system has an ongoing role in this process, we combined electrophysiological, pharmacological and optogenetic techniques in awake head-fixed mice, as they were presented with auditory stimuli passively. We discovered that PFC responds to sensory input under these conditions, and that cholinergic signaling in the AC plays an important role in modulating the strength of PFC responses through muscarinic receptor activation. These findings suggest that cholinergic interactions are routinely involved in PFC sensory gating thereby contributing to normal sound processing.

## 2. Materials and Methods

All procedures involving animals were approved by the Boston University Institutional Animal Care and Use Committee (IACUC). A total of 38 transgenic mice were used in this study (3-6 months-old, on the day of recording).). 32 Chat-ChR2 transgenic mice were obtained by crossing Chat-Cre mice (B6;129S6-Chat^tm1(cre)Lowl^/J) with Ai32 mice (B6;129S-Gt(ROSA)26Sor^tm32(CAG-COP4*H134R/EYFP)Hze^/J (both were obtained from Jackson Laboratory, Maine). 6 control Ai32 animals served as controls for optogenetic experiments.

### 2.1 Surgical Procedures

Mice were surgically implanted with a head-plate as described previously (McCarthy et al., 2011). Briefly, under isoflurane anesthesia, a custom head-plate designed to allow access to PFC and AC was anchored to the skull with 3 stainless steel screws. A fourth screw was connected to a metal pin and placed in the skull above contralateral parietal cortex to serve as the ground.

### 2.2 In Vivo Electrophysiology

Upon complete recovery from surgery, typically one week, mice were first habituated to the head-fixed condition in the recording apparatus, enclosed in a custom sound attenuation chamber (Industrial Acoustics Company Inc., Bronx, NY) the day before the recording experiment was performed. During the recording and habituation sessions, mice were head-fixed to a custom holder that was anchored to the recording table, and loosely wrapped in a breathable mesh (Butler Schein, Dublin, OH; Vet-Flex EZTear). On the recording day, electrode probes were slowly lowered through two small craniotomies into the PFC (AP+2.0, ML +0.4, DV −2.75) and the AC (AP −2.3 to +3.6, ML +4.0 to +4.5, DV −1.0) using motorized micromanipulators (Siskiyou, Grants Pass, OR) at 100-200 µm/min. A 16 contact linear probe (Neuronexus, Ann Arbor, MI; model : A1×16-10mm-100-177-A16) with 100 um spacing between electrode contacts was inserted dorsal-ventrally into the PFC. A 32 channel probe (4 shanks, 8 sites per shank with 100 um spacing between electrode contacts and 400 um spacing between shanks, Neuronexus, Ann Arbor, MI; model: A4×8-5mm-100-400-177-A32) was positioned into the AC, perpendicular to the cortical surface. Because of the curvature of the AC cortical surface, not all four shanks could be placed at precisely the same depth during each experiment. Probes were advanced until all electrode contacts were within the cortical tissue and shanks were positioned along the rostro-caudal axis of the AC. An optical fiber, 200 um in diameter, was lowered through a third craniotomy using a microdrive (Siskiyou, Grants Pass, OR) into the nucleus basalis (NB; AP +0.5, ML +1.5, DV −4.5) for optogenetic stimulation.

Extracellular recordings were made with a TDT multichannel RZ2 recording system (Tucker Davis Technologies, Alachua, FL). To record spikes, signals were digitized at 24414 Hz and band pass filtered between 300-5000 Hz. Spikes were identified by threshold crossings, and the thresholds were manually set at the beginning of each recording session. Timestamps and waveform snippets (1.3 ms long) were stored for later offline processing. Local field potentials (LFPs) were low pass filtered at 1000 Hz, and digitized at 3051.8Hz.

### 2.3 Pharmacology

The 32 channel recording probe in AC was also coupled to an infusion pipette under the control of a picospritzer for local drug infusion (Figure 1B). The infusion pipette was a pulled glass pipet with an opening of approximately 5 µm in diameter, and was placed so that the tip was halfway between the two innermost shanks and terminated halfway between the deepest and most superficial electrode contacts. After a baseline recording session, a picospritzer was used to deliver 500 nL of drug or vehicle into the AC over 5-10 minutes. Post-drug recording sessions were performed 20-30 minutes after the completion of drug infusion to allow for diffusion. We infused the muscarinic antagonist scopolamine at a low dose of 10ug/uL and a high dose of 100ug/uL, and the nicotinic antagonist mecamylamine at a dose of 1ug/uL. All drugs were dissolved in artificial cerebro-spinal fluid (ACSF, Tocris Bioscience, Bristol, UK). ACSF was used as control for infusion experiments.

**Figure 1.**
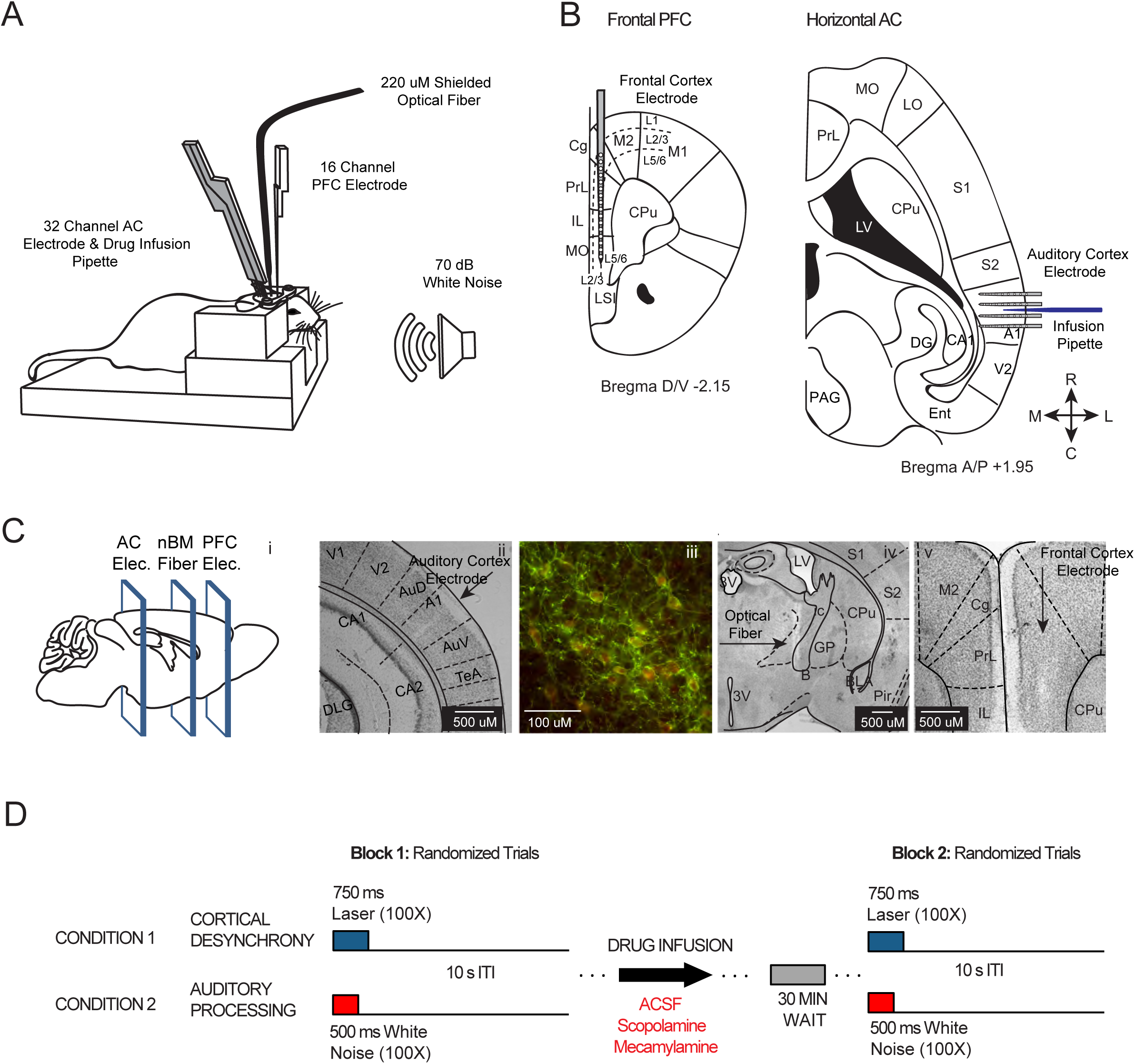
(Color): Experimental paradigm. **A:** Head fixed mouse preparation with electrode and optical fiber placements during recording. One 16-channel electrode was positioned in PFC, an infusion pipette positioned between the two inner shanks of a 32 channel, 4 shank recording electrode in AC, and a shielded optical fiber in nucleus basalis (NB). **B:** Anatomical location of recording electrodes and infusion pipette represented in a frontal (PFC) and horizontal section (AC). PFC single shank penetration occurred along the dorsal-ventral axis with sites facing medially. For the AC array, insertion was at a 30° angle with all recording sites facing medially while the shanks and infuser were situated along the rostral-caudal axis. **C (i):** Cartoon depiction of electrode and fiber placement for histology sections shown to the right. **(ii):** Section from AC showing electrode placement for this recording area. **(iii):** Histology image showing ChR2-GFP expression in tissue co-labeled with antibodies directed against choline acetyl transferase (Chat). Note, ChR2-GFP expression is limited to cells also expressing Chat (red). **(iv):** Nissl stain showing optical fiber track and placement in NB**. (v):** Section from frontal cortex demonstrating electrode placement from a representative animal. **D:** During the preinfusion recording period (block 1), mice were presented with 100 trials of optical stimulation alone and 100 trials of auditory stimulation alone, with the order of trials shuffled throughout the block. A second recording session was performed 30 min after drug infusion (block 2). **Abbreviations:** Cg; Cingulate Cortex, Prl; Prelimbic cortex, IL; Infralimbic Cortex, LSI; CPu; Caudate Putamen, MO; Medial Orbital Cortex, LO; Lateral Orbital Cortex, S1; Primary Somatosensory Cortex, LV; Lateral Ventricle, AuD; Secondary Auditory Cortex Dorsal, A1; Primary Auditory Cortex, AC; Auditory Cortex, PFC; Prefrontal Cortex, nBM; Nucleus Basalis of Meynert, B; Basal Nucleus of Meynert, 3V; Third Ventricle; R; Rostral, C; Caudal, M; Medial, L; Lateral).

### 2.4 Auditory Stimulus

White noise auditory stimuli were generated with a RZ2 Bioamp processor and RP2.1 real time processor (Tucker Davis Technologies), and digitized at a frequency of 48828 kHz. Stimuli were played with a multi-field magnetic speaker (Tucker Davis Technologies, model: MF1), positioned at 20 cm from the animal’s head, and calibrated to be 70 dB sound pressure level (SPL) with a conditioning amplifier and microphone (Bruel and Kjaer, Naerum, Denmark; amplifier: model: type 2690—0S2; microphone: type 4939-A-011). The stimuli consisted of 500 milliseconds white noise bursts (5ms linear ramp to 90% of peak amplitude).

### 2.5 Optogenetic Stimulation

We positioned an optical fiber in the nucleus basalis (NB) in order to optogenetically stimulate cortically projecting cholinergic neurons in Chat-ChR2 transgenic mice that express ChR2 selectively in cholinergic neurons, while simultaneously recording in AC and PFC (Figure 1D, condition 1). Laser light for optogenetic stimulation of NB was delivered through a multimode fiber, 200 um in diameter, optically shielded (Thorlabs, Newton NJ; model: BFH48-200), and coupled to a 473nm DPSS laser (Shangai Laser Ltd., Shanghai, China; model: BL473T5-200FC). Laser power was calibrated to 15mW at the fiber tip prior to insertion. 40 Hz square light pulses (20% duty cycle) lasting 750 ms were delivered via TTL trigger from the RZ2 recording system.

### 2.6 Recording Session

The recording protocol consisted of combinations of optogenetic and auditory stimuli. Two trial types were analyzed, one consisted of a sound burst alone, and the other consisted of a 750 ms pulse of light delivered alone. Both trial types were presented to mice 100 times, with their order randomized. The inter-trial interval (ITI) was 10 seconds.

### 2.7 Histology

At the end of the experiments, all mice were anesthetized and transcardially perfused for histological verification of electrode placement using cresyl acetate (C-1893; SigmaAldrich, Natick MA). Mice were perfused with 30 mL buffered saline, followed by 30 mL 4% paraformaldehyde. Brains were carefully removed and post-fixed overnight in 4% paraformaldehyde before being transferred to a 30% sucrose solution, and then sectioned coronally in 40µm slices with a freezing microtome (CM 2000R; Leica). Tissue sections were collected throughout the basal forebrain, AC, and PFC. Alternate sections were then rinsed with 0.05M PBS buffer and mounted on gelatin-coated slides and allowed to dry overnight. Sections were then rehydrated by a descending series of alcohol rinses (2 min each in 100%, 95%, 90%, 70%, and 50%) before being placed in deionized water for 5 min. Following rehydration, sections were incubated in 0.1% cresyl acetate for 5 min, followed by dehydration in an ascending series of alcohol rinses (50%, 70%, 90%, 95%, 100% (2X) for 3 min each and cleared with xylene (534056; SigmaAldrich, Natick MA) for 15 minutes. Slides were then coverslipped using DPX (06522; SigmaAldrich, Natick MA) mounting medium and analyzed.

To validate ChR2 expression patterns, we also performed immunohistochemistry with antibodies directed against ChAT (Choline acetyltransferase is selectively expressed in cholinergic neurons) as described previously (Deckmann et al., 2014). Briefly, sections were rinsed with 0.05 M Tris-HCL-buffer (Tris, pH=7.6), followed by 60 mins rinse in blocking buffer containing 5% serum (Jackson ImmunoResearch, 017-000-121) and 0.2% Triton. Sections were incubated for 24 hours with goat anti-ChAT antibody (AB144P, Millipore; Temecula, CA) diluted at 1:500. Sections were then rinsed 3X at 10 min each in Tris-HCL and then incubated with the secondary antibody Alexa Fluor 594 (donkey-anti-goat; Life Technologies A11058; Grand Island, NY) at 1:200 for 2 hours. Sections were rinsed again and then mounted on gelatin-coated slides using anti-fade mounting media Vectashield (H-1400, Burlingame, CA).

### 2.8 Data Analysis

All data analysis was performed with custom Matlab functions (MathWorks, Natick, MA). Statistical tests were Friedman’s test or ANOVAs, for comparisons of drug and proximity to infusion pipette, Wilcoxon signed-rank tests and paired t-tests for comparisons between baseline and post-infusion measurements, and Pearson’s linear correlation to compute correlation between response magnitude and recording depth. In some instances the logarithm of the data values were used for statistical analysis.

LFPs were first bandpass filtered between 1 Hz and 150 Hz, and then down-sampled by a factor of 8 (381 Hz) prior to further analysis. In awake head fixed conditions, we occasionally found motion induced artifact. To remove trials contaminated with motion artifact from subsequent analysis, we took the root mean square (RMS) in 100 ms bins for each trial, and rejected trials if the averaged RMS exceeded 5 standard deviations at any time throughout a trial.

### 2.9 LFP spectral analysis

We used the open source software package Chronux (www.chronux.org) to compute multi-taper spectrograms, using three tapers and a 500 ms sliding window with 5 ms step. Single trial spectrograms were first calculated, and then averaged across trials. To examine the effects of optogenetic stimulation, we calculated normalized spectrograms by dividing the average power during each time window by the mean power during baseline period, defined as 5 second window immediately preceding optogenetic stimulation. Time-frequency windows of interest were quantified by summing all normalized power within given frequency and time ranges.

To quantify the influence of pharmacological agents on baseline LFP power for comparison between the baseline and post-drug infusion condition, we calculated multi-tapered power spectra during the 4-9 second window of the ITI. Note, the ITI is 10 seconds long (Fig. 1). We calculated the log(power) at 10-100 Hz and 1-10 Hz frequency bands, and compared the log(power)before and after drug infusion. We also computed cortical desynchrony by taking the ratio of power at 10-100Hz and power at 1-10Hz. To examine the effect of optogenetic stimulation of NB on cortical desynchrony, we compared cortical desynchrony during the 750ms light stimulation window to the 750 ms time period immediately before optogenetic light illumination onset as the baseline.

### 2.10 Current source density (CSD) analysis

CSD analysis estimates the second spatial derivative of LFP signals to determine the relative current across the cortical laminar depth. CSDs were calculated using LFPs recorded from the four laminar probes inserted in AC, as described previously (Mitzdorf, 1985; Nicholson and Freeman, 1975). We averaged LFPs recorded from each electrode contact across all trials prior to CSD calculation.

First, we applied spatial smoothing across the eight electrode contacts within the same shank as described in (Sakata and Harris, 2009).

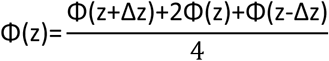

Where z is the depth perpendicular to the cortical surface, Δz is the electrode spacing, and Φ is the potential. Then we estimated the CSD as described in (Mitzdorf, 1985; Nicholson and Freeman, 1975).

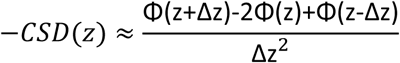

To quantify the total current across cortical lamina during temporal windows of interest, we averaged the root mean square (RMS) of CSD curves at all depths. Additionally, for population analysis of CSDs between drug-treatments, we separately averaged CSD RMS values from either the two inner shanks or the two outer shanks, to reflect the spatial symmetrical orientation of the probe shanks with respect to the infusion pipette (Figure 1B, right, 4A). For display purposes, CSDs were interpolated linearly and plotted as pseudocolor images.

We analyzed the CSD latencies for both the current sink onset and the current source peak across drug conditions to determine the effects of drug infusion on AC sound evoked responses. We first identified the electrode contact on each laminar shank with the largest current sink, calculated as the average amplitude within 50 ms of sound onset. This electrode contact was used to quantify CSD onset latency. Sound onset latency was chosen as the point at which the CSD crossed 3 S.D. from baseline. The end of the sink was calculated by taking the time point at which the rising phase of the CSD crossed 3 S.D. from baseline. CSD profiles were not used to estimate current in PFC, because the probe was not oriented perpendicular to the cortical lamina.

CSD analysis was also used to determine the granular layer on each shank in AC. Along each shank, the channel that had the earliest sink was considered to be in the granular layer. This layer identification was used to select granular layer channels for ERP latency, and used for anatomical classification of laminar CSD magnitudes and MUA firing rates. For laminar analysis of CSD curves, peak-trough values were taken from the mean supragranular and infragranular channels, as well as granular channels, for each mouse. Additionally, CSD traces were used for spectral analysis of the baseline LFP. When used for spectral analysis—in which averaging occurs in the spectral domain—spatial smoothing was not applied to the CSD.

### 2.11 Event related potentials (ERPs) analysis

To quantify auditory ERPs in PFC, we averaged LFPs recorded across all trials, and all electrode contacts of the 16 channel linear probe in PFC. ERP magnitude was calculated by subtracting the minimum from the maximum value of the ERP within 100 ms following stimulus onset. In PFC, ERPs at various depths were either averaged together, or analyzed separately to examine depth-dependence. ERP latencies were chosen as the point at which the ERP exceeded 3 S.D. from baseline. We limited our analysis of latency to trial-averaged ERPs that exceeded 3 S.D. of the baseline activity within 100 ms post sound onset in PFC (28 of 32 recordings).

To facilitate latency comparisons between PFC and AC, we also calculated AC latencies using ERPs. ERPs were taken from the electrode contact centered on the granular layers and ERPs that did not exceed 3 S.D. in this window were excluded. Latency values were then calculated from AC granular layer ERPs in the same manner as PFC latencies. In instances where a latency values were less than 10ms, those events were not included (recordings from 2 out of 128 total shanks) as it is physiologically improbable to be a result of auditory input and more likely to be a result of random deviation in the baseline LFP.

### 2.12 Spike analysis

Spike snippets were detected with manual thresholds during recording sessions and analyzed offline. Multi-unit (MU) activity was reported consistent with other auditory studies using laminar electrodes (Guo et al., 2012; Steinschneider et al., 2008). This was done because of generally lower signal amplitude obtained with laminar probes and the inability for each electrode contact to be independently positioned to best capture nearby single neuron activity leading to inconsistencies in single-unit isolation (Bragin et al., 2000).

We calculated the width of spike waveforms at half of their peak-trough maximum. Spike events were only included if their width at half-maximum was between 0.1 and 0.4ms. Additionally, coincident events indicative of motion induced artifact across multiple contacts on the same probe were removed if they occurred within two samples of any other spike (83 µsec).

MUA peri-stimulus time histograms (PSTHs) were calculated using 2 ms time bins for population and statistical analysis. AC MUAs were considered sound responsive if the mean z-score exceeded an absolute value of 2 at early onset (10-30 ms) or sustained an average z-score of 1 throughout the duration of the 500 ms auditory stimulus. Similarly, PFC MUAs were considered responsive if their mean z-score exceeded an absolute value of 2 during the 10-60 ms window after sound onset or exceeded 1 throughout the duration of the stimulus. Firing rate curves were binned with a 16 ms moving average for display purposes. In AC, in order to measure the firing rate (FR) decline following sound onset, we measured the integrated firing rate for each MU from its largest peak in the first 50ms of sound onset until 200 ms post sound onset (Figure 6A, 6C).

The peak evoked PFC activity was calculated as the maximum of the mean population firing rate across all MUAs. Peak PFC firing rates were calculated for each population defined by anatomy and drug group individually. Cross-correlograms for AC MUAs within the duration of the sound window (0-500ms) were calculated between pairs of electrode shanks and across contacts within each shank. Cross-correlograms were normalized by the mean number of spike events per bin multiplied by the length of the cross-correlogram. To compare latencies of MUAs in PFC and AC, MUs with early burst-type response patterns were selected and the latencies to the half-maximum of their firing rate curves were used to estimate response latencies. Significantly more early burst-type MUs were present in AC (492 MUs) than in PFC (40 MUs).

## 3. Results

### 3.1 Prefrontal cortex (PFC) shows subregional variations in auditory evoked responses that lag responses in auditory cortex (AC) during passive sound processing

While it has been well established that neurons in PFC respond robustly to auditory stimuli that are relevant to behavioral tasks (Chang et al., 2010; Euston et al., 2012; Hyman et al., 2012; Maren and Quirk, 2004; Moorman and Aston-Jones, 2015), few studies have explored the extent to which subregions of PFC are responsive to the passive presentation of sound in awake rodents. A small number of studies have previously identified PFC neural activity in response to auditory stimuli − CS presentations before fear conditioning for example (Burgos-Robles et al., 2009; Chang et al., 2010), but none to our knowledge, have compared the response properties between AC and multiple regions of PFC simultaneously in awake animals. In order to better understand how sound is coincidently represented in AC and PFC during passive sound processing, we recorded from both structures using multi-contact electrode probes (Figure 1A-C) in awake-head fixed mice. Animals underwent acclimation and habituation to the recording configuration and sound booth prior to the recording session. This experimental design is similar to other experiments where naturalistic stimuli were delivered to habituated head fixed animals (Hromadka et al., 2008; Lee et al., 2012; Pinto et al., 2013). Acoustic white noise was used as a stimulus to activate as much of the AC tonotopy as possible, in order to evaluate the strength of interactions across regions. To reduce potential for stimuli to have an aversive quality, we presented the stimulus at an intensity of 70dB sound pressure level (SPL). Our goal was to capture excitatory and inhibitory interactions across multiple tonotopic regions during a single stimulus so that we could test the effects of cholinergic antagonists on auditory responses.

We first quantified the multi-unit activity (MUA) recorded in PFC, and found that the 12.0% of multi-units (MUs) significantly increased their firing rate during sound presentation, while 0.2% decreased (n=502 MUs recorded in 32 mice; Figure 2A, C). These findings are similar to a number of studies that assessed responsive neurons (∼11-25%) in prefrontal areas for conditioned auditory stimuli prior to fear conditioning (Burgos-Robles et al., 2009; Chang et al., 2010). We also observed that MUs exhibited complex temporal dynamics, showing a combination of monophasic and ramped firing responses upon sound presentation. Responses were generally followed by a slight reduction after sound offset that lasted for ∼500 ms before returning to baseline (Figure 2C). The observed response profile as well as the mean latency of the responses were similar to those noted in ventral orbital cortex in head restrained mice using a white noise auditory stimulus (Winkowski et al., 2017). Because the mouse PFC is layered medial-laterally and is parallel to the dorsal-ventral axis of electrode insertion (Figure 1B, Figure S7), we could not sample across the cortical layers as we could with AC. Instead, the PFC linear electrode targeted the middle cortical layers (deeper layer three and superficial layer five) of a number of anatomically dissociable PFC sub-regions, including the cingulate (Cg), prelimbic (PrL), infralimbic (IL), and medial orbital (MO) cortices. Supplemental Figure 7 shows the electrode penetrations, in relationship to the anatomical boundaries of PFC. We found that while all PFC sub-regions were responsive to the auditory stimulus, they differ in their response magnitude, with IL being less responsive than the more superficial Cg, PrL, or deeper MO subregions (chi-square=6.43, p=0.01; Figure 2E).

**Figure 2.**
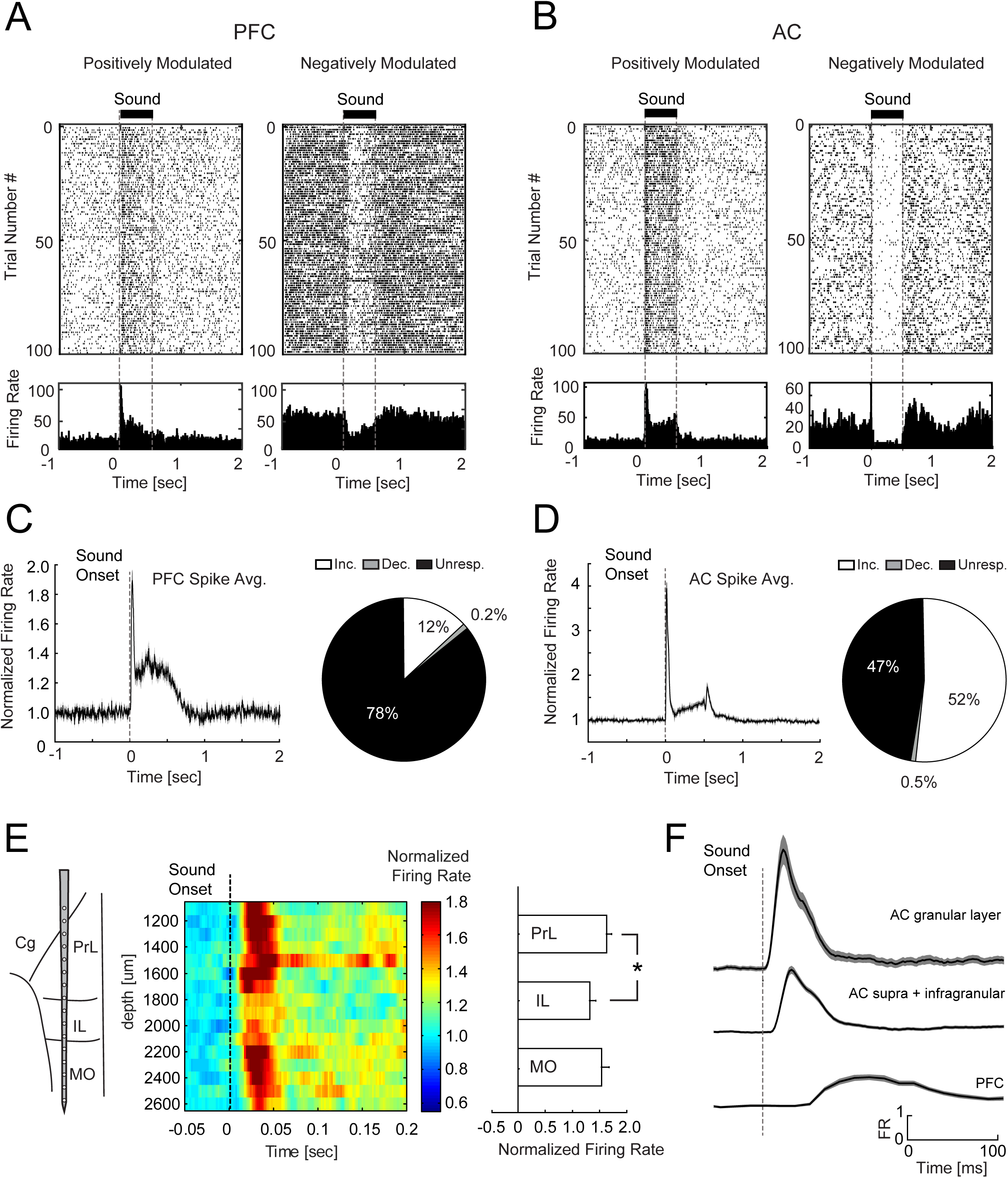
(Color): Auditory stimuli evoked strong MUA in PFC and AC during passive sound presentation in mice. **A:** Example MU from PFC that shows an increase (left) or a decrease (right) in firing rate during the sound stimulus. **B:** Example MU from AC that shows an increase (left) or a decrease (right) in firing rate during the sound stimulus. **C, D:** Mean population MUA in PFC (**C**) and AC (**D**) during the sound presentation (left) and the proportion of responsive MUs for each region (right). **E:** Mean normalized PFC MUA from all animals across all depth locations from the pre-infusion condition. Note that all regions show an increase in firing rate with prelimbic (PrL) exhibiting the largest increase in firing rate (* = p<0.05). **F:** Mean population MUA from granular and extragranular layers in AC and mean population MUA from the PFC from the pre-infusion period. (Cg; Cingulate Cortex, Prl; Prelimbic cortex, IL; Infralimbic Cortex, MO; Medial Orbital Cortex).

In contrast, the majority of AC MUs showed sound modulation: 52% of MUs increased their firing rate during sound presentation, and 0.5% decreased (n=980 MUs recorded in 32 mice; Figure 2D, right). The neurons with suppressed firing rates showed higher baseline firing rates when compared to neurons that increased their firing rate (baseline: 42.24±8.65 Hz; mean±SEM: suppressed group. vs. 35.97±6.92 Hz; mean±SEM activated group). These findings are consistent with other studies demonstrating that broadband auditory stimuli evoke a relatively sparse, yet tonotopically extensive response in awake, un-anaesthetized animals (Evans and Whitfield, 1964; Hromadka et al., 2008). MUs typically exhibited a large transient increase immediately following sound onset, but some channels also showed low levels of sustained activation throughout the duration of the stimulus (Figure 2B, left). We found that MUs granular layers (based on current source density (CSD) profiles – described below), showed the greatest increases in firing rates at sound onset, while the remaining sites showed more gradual responses that were broader in duration (Figure 2F). MU activity also occurred significantly earlier in AC compared to prefrontal cortex (t(5,30)=-2.132; p=0.034; Figure 2F).

We further examined the auditory stimulus evoked local field potentials (LFPs) recorded in both AC and PFC and found large event related potentials (ERPs) on all recording sites in PFC and AC (Figure 3A). PFC ERPs had an average onset latency of 28.48±7.39 ms (mean±S.D., n=32 mice, Figure 3Bi-3Di). Interestingly, we detected a significant correlation between ERP amplitude and dorsal-ventral depth (R=-0.154, p= 5e-4, n=512 recording sites, 32 mice), with the largest ERPs observed on the most superficial subregions (Cg and PrL) where the firing rate increases were also greatest (Figure 2E). We also detected a systematic increase in latency with depth, with the deepest recording sites lagging the most superficial recording sites by ∼5 ms (Figure 3Ci). The changes in ERP amplitude and latency along the dorsal-ventral axis of PFC suggest that mouse PFC subregions receive differing levels of auditory input, with the superficial subregions (i.e., Cg and PrL) receiving auditory input earlier and of greater magnitude than deeper subregions (i.e. IL and MO).

**Figure 3:**
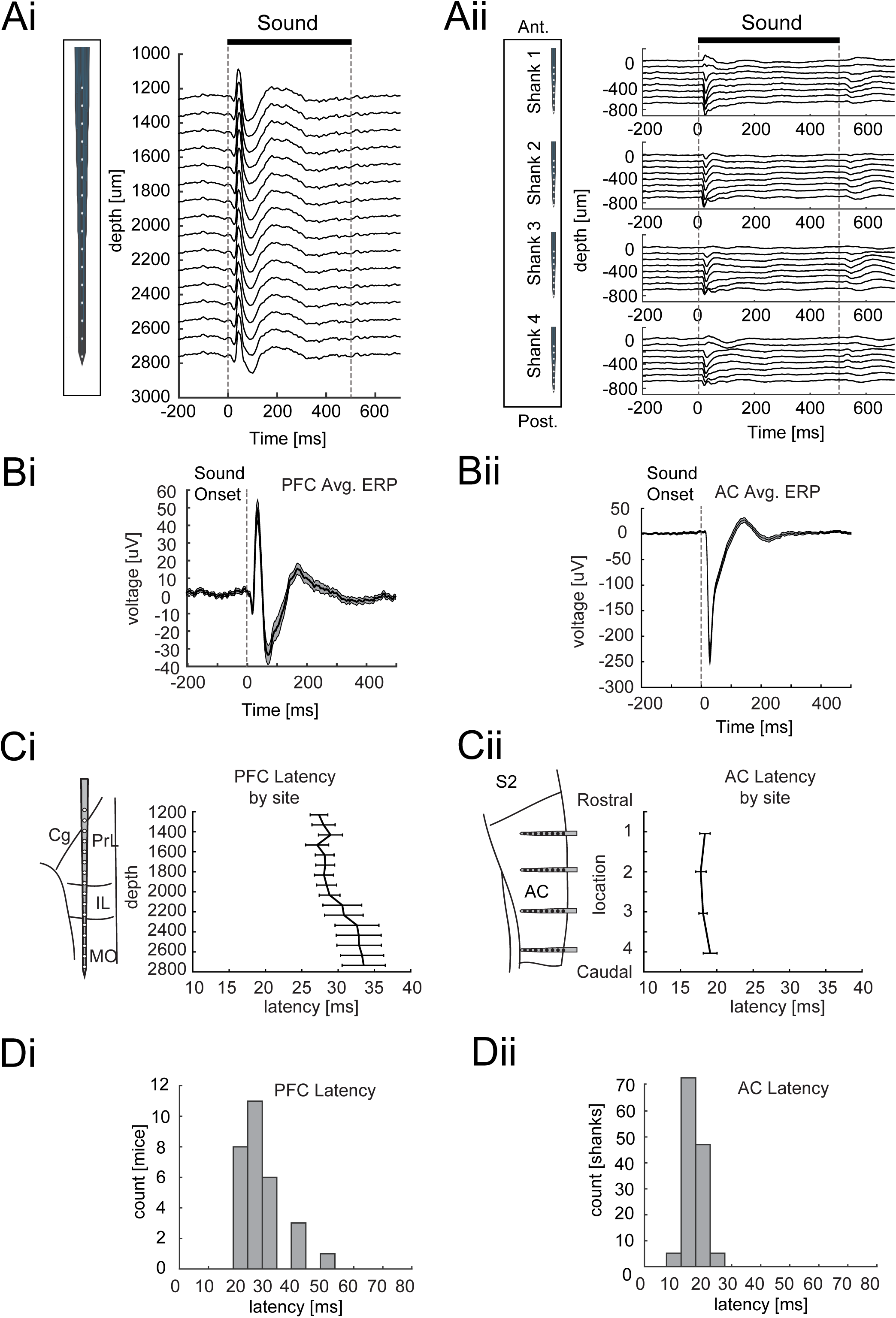
Auditory input produced robust ERPs in PFC and AC during passive sound presentation in mice. **A:** Representative ERP response to auditory stimuli across all channels in the PFC (i) and AC (ii). **B:** Population ERP from PFC (i) and AC (ii), in response to auditory stimuli. **C:** ERP onset latency along the dorsal-ventral axis of PFC (i) and along the rostro-caudal axis of AC (ii). (solid lines are mean, and shaded areas are mean ± s.e.m. **D:** Histogram of ERP onset latencies from PFC (i) and the granular layers of AC (ii).

In AC, robust ERPs were evoked by auditory input at most recording sites, with some sites exhibiting larger ERPs than others, presumably because some sites were within regions of the AC tonotopy that were outside the frequency range of the band-limited white noise auditory stimulus used (sampled at 48.8 kHz). Across the cortical depth, we found that the largest ERPs were in the middle layers, usually the two middle electrode sites, ∼300-500 um from the surface of the cortex (Figure 3Aii). AC ERPs paralleled the evoked MUAs, with the most robust responses observed immediately after sound onset (Figure 3Bii, 2D), consistent with previous studies investigating AC (AI, AAF, or AII subregions) (Joachimsthaler et al., 2014; Stiebler et al., 1997). The mean onset latency for ERPs in AC granular layers (defined by current source density analysis; see methods) was 17.89±2.95 ms (mean±S.D., n=126 shanks, 32 mice; Figure 3Cii, Dii). When we compared the ERP response latencies, we found that activity also occurred significantly earlier in AC compared to prefrontal cortex (Wilcoxon rank-sum test, p=1.8e-11), consistent with the MU latencies described earlier. The difference of ∼10ms in activation between PFC and AC is consistent with that observed in ketamine anesthetized rats where a 8-18ms delay was observed between A1 and PFC (Martin-Cortecero and Nunez, 2016). Taken together, these results demonstrate that mouse PFC is activated by auditory stimuli during passive sound processing, and superficial PFC sub-regions receive stronger and earlier auditory input.

### 3.2 Local muscarinic receptor antagonism disrupts cortical desynchrony in AC by altering intracortical connectivity

We configured our AC recording probe with a microinfusion pipette that was positioned between the two inner shanks of the four shank electrode in AC for targeted drug delivery (Figure 1B, 4A). The nicotinic blocker, mecamylamine (0.5µg/500nL), was delivered at a dosage shown to be effective in altering sound evoked response latency and characteristic frequency thresholds when locally delivered into AI in rodents (Liang et al., 2006; Metherate, 2004). However, we could not find similar studies in which the muscarinic blocker scopolamine was delivered intracortically into AC during sound processing. Therefore we chose a high dose (50µg/500nL) and a low dose (5 µg/500nL) that border the broad range of effective doses cited in the literature as influencing behavior in active and passive experimental paradigms, as well as disrupting the influence of cholinergic activation in cortical circuits (Boix-Trelis et al., 2007; Herrero et al., 2008; Ingles et al., 1993; Maruki et al., 2003; Rogers and Kesner, 2004; Santucci and Shaw, 2003; Zhou et al., 2011).

**Figure 4.**
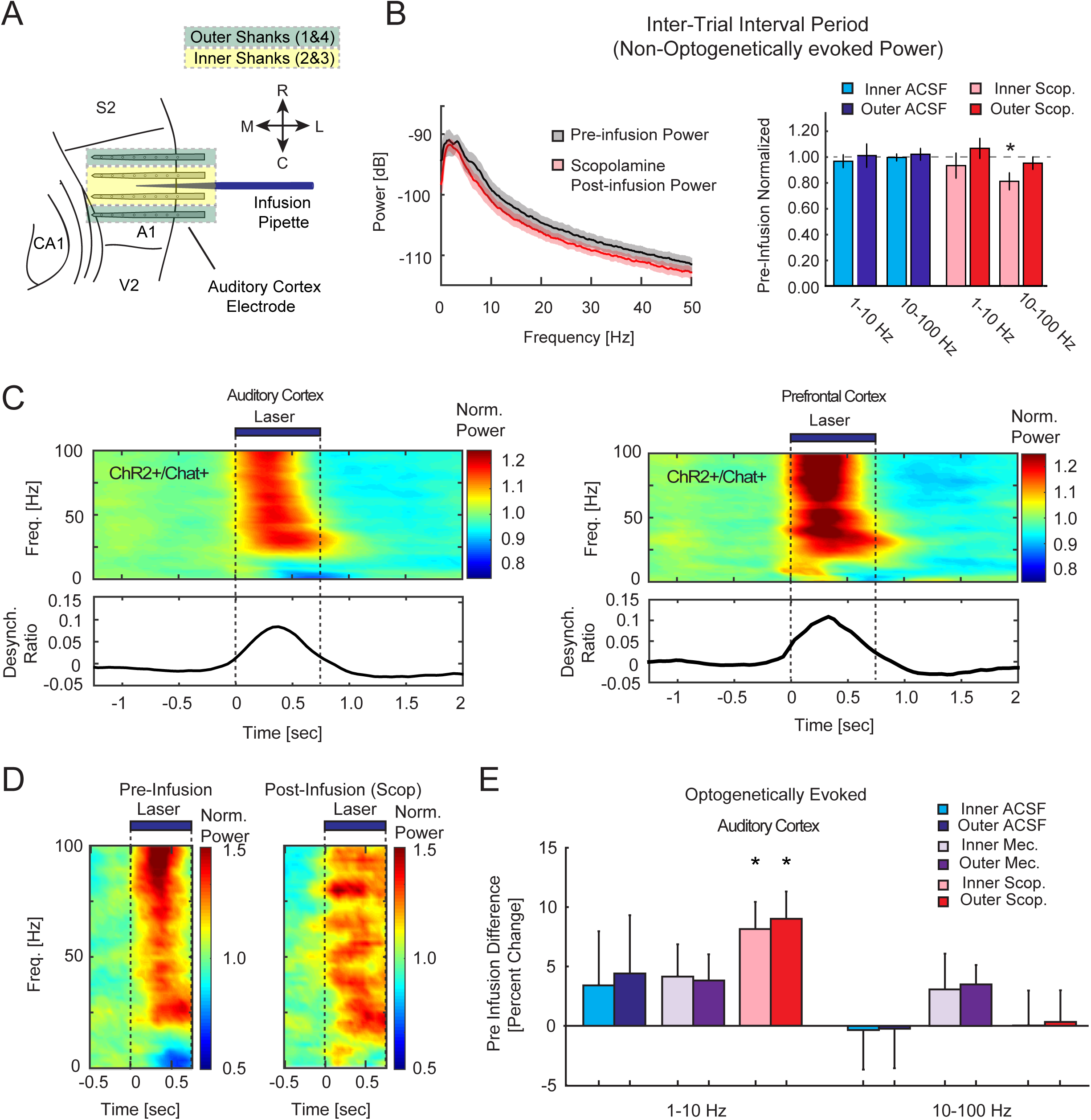
(Color): Effects of cholinergic antagonism on LFP power and optogenetically induced cortical desynchrony in AC and PFC. **A:** Diagram illustrating recording/infusion configuration in AC in a horizontal section. Compass denotes rotral-caudal and medial-lateral axis. Inner shanks are highlighted yellow and outer shanks are highlighted green. Pipette location is noted in blue. **B:** Population spectrum of LFP power on inner shanks in AC before (black) and after (red) infusion of high dose scopolamine during the inter-trial interval in the absence of stimuli or optogenetic activation of cholinergic NB (left). Bar height represents population mean ± s.e.m normalized to pre-infusion baseline across all mice recorded (right) and compares ACSF (blue) to scopolamine (red). Outer sites are darkly shaded relative to inner sites (*= p<0.05). **C:** Cortical desynchrony produced by optogenetic activation of cholinergic NB. Trial averaged, baseline normalized spectrograms from AC (left) or PFC (right) during optical stimulation. The ratio of low frequency power (1-10 Hz) to high frequency power (10-100 Hz) for AC and PFC is shown below (bottom). **D:** Population spectrogram of AC LFP during optogenetic activation of cholinergic NB prior to high dose scopolamine infusion (left), and following infusion (right). E: Change in AC LFP power by frequency range during optogenetic stimulation in the presence of antagonists or vehicle. Values are expressed as percentages of pre infusion power (mean ± s.e.m). Outer sites are darkly shaded relative to inner sites (*= p<0.05).

As systemic application of cholinergic antagonists have been shown to modulate the cortical LFP power spectrum (Kalmbach and Waters, 2014), we first examined the effects of cholinergic antagonism on LFP power spectra during the inter-trial interval (ITI) period relative to the pre-infusion ITI period (Figure 4B). Neither ACSF, mecamylamine, nor low dose scopolamine altered the LFP power spectrum, on inner or outer shank locations following drug infusion (Wilcoxon signed-rank test, p>0.05). However, infusion of high dose scopolamine reduced LFP power at higher frequencies (10-100Hz) on the inner electrode shanks closest to the infusion site (Wilcoxon signed-rank test, p=0.02, n=9 mice; Figure 4B), but not on the outer shanks distal to the infusion site (Wilcoxon signed-rank test, p>0.05). This effect confirms that 500 nL scopolamine delivery was largely isolated to the immediate vicinity of the infusion site. We found no effect of scopolamine infusion on low frequency power on either the inner or outer shanks (1-10 Hz: all Wilcoxon signed-rank test, p>0.05, n=9 mice; Figure 4B). Because field potentials can express volume conduction, particularly at low frequencies, we wanted to be certain that scopolamine did not induce reductions in LFP power that may have been masked by volume conductance into our recording area from adjacent regions away from the infusion site. Therefore, we further analyzed CSD waveforms, the second spatial derivatives of LFPs that estimate local current, and found no significant change in power in the 1-10Hz range on inner or outer shanks (all Wilcoxon signed-rank test, p>0.05, n=9 mice; Figure S10). Analysis of the CSD waveform also revealed that only the inner shanks in the 10-100Hz frequency range showed significant reductions following scopolamine infusion (Wilcoxon signed-rank test, p=0.02, n=9 mice), recapitulating the only significant effect for the baseline LFP analysis.

We then examined the efficacy of cholinergic antagonism on cortical desynchrony, defined as the ratio of LFP power at high frequency (10-100 Hz) to low frequency (1-10 Hz). Cortical desynchrony is a widely observed phenomenon that coincides with cholinergic activation of sensory cortex, and is characterized by a simultaneous decrease of LFP power at low frequencies (<10 Hz) and an increase in power at high frequencies (10-100 Hz) (Goard and Dan, 2009; Kalmbach et al., 2012; Kalmbach and Waters, 2014; Metherate and Ashe, 1993; Pinto et al., 2013). We induced cortical desynchrony through optogenetic stimulation of nucleus basalis (NB) in Chat-ChR2 transgenic mice selectively expressing ChR2 in NB cholinergic neurons (Figure 1C). Prior to drug infusion, we found that optogenetic stimulation of NB cholinergic neurons induced cortical desynchrony in both AC and PFC (Wilcoxon signed rank test, p=1.405e-6 for PFC; p=6.48e-7 for AC, n=32 mice). Cortical desynchrony in AC and PFC was short lasting, limited to the light stimulation window, and was not detectable in control Ai32 transgenic mice (Supplemental Figure 1). Infusion of high dose scopolamine reduced cortical desynchrony, selectively blocking the decreases in low frequency power (chi-square=15.8, p= 5.16 e-6, inner and outer shanks) on both inner and outer shanks in AC, but not the increases in high frequency power (Wilcoxon signed-rank test, p>0.05), suggesting that only the low frequency component of cortical desynchrony depends on mAChR activation. Neither low frequency nor high frequency power was impacted by ACSF vehicle, mecamylamine, or low dose scopolamine infusion (Figure 4E; Wilcoxon signed-rank test, all p>0.05). Disruption of low frequency components of desynchronization is consistent with previous studies (Kalmbach and Waters, 2014; Metherate et al., 1992), and confirmed that high dose scopolamine was effective in blocking muscarinic receptors and neutralizing the influence of intrinsic cholinergic activation.

It is interesting that the muscarinic blocker scopolamine reduced cholinergically induced cortical desynchrony across broad cortical areas, recorded on both the outer and the inner recording shank locations (Figure 4E). Since scopolamine only altered LFP power during the ITI at inner shank locations, and not outer shanks locations (Figure 4B), the observed effects of scopolamine on cortical desynchrony on the outer shanks cannot be explained by direct antagonism of muscarinic receptors at the outer shanks. Thus, scopolamine’s effect on cortical desynchrony at the outer shanks must be mediated by network interactions between neurons at the inner shank locations and neurons at the outer shank locations. This result suggests that acetylcholine, through activating mAChRs, modulate lateral interactions between neurons of adjacent regions of AC, which may contribute to the tonotopic refinement of auditory input. Together, these results demonstrate that acetylcholine induced cortical desynchrony contributes to the interactions of cortical networks through influences on lateral connectivity.

Optogenetic activation of NB also produced robust cortical desynchrony in PFC that was not blocked by vehicle or any of the antagonists infused into AC (Supplemental Figure 5; all Wilcoxon signed-rank test, p>0.05). Many previous studies have linked cholinergic activation to cortical desynchrony in sensory or motor cortices. Our observation that cortical desynchrony is also present in PFC, highlights the broad projection patterns of basal forebrain cholinergic neurons (Zaborszky et al., 2012). While high dose scopolamine infusion in AC altered cortical desynchrony in AC, it failed to alter cortical desynchrony in the PFC suggesting the effects produced by local scopolamine infusion in AC were confined to AC, and not relayed to PFC.

### 3.3 Optogenetic stimulation did not alter sound evoked amplitude

Because previous work has demonstrated that behavioral training or basal forebrain (BF) stimulation can significantly reorganize receptive fields or temporal response profiles in AC (Fritz et al., 2007; Froemke et al., 2007; Kilgard and Merzenich, 1998a, b; Polley et al., 2006), we wanted to be certain that our optogenetic stimulation parameters did not promote plasticity changes during recording. Traditionally this plasticity occurs over days or weeks and often involves stimulus-response based learning with coincident activity, although rapid, within session plasticity has been reported (Froemke et al., 2007). Plasticity measures often involve changes in the tuning of AC neurons modulated by context with behaviorally relevant auditory stimuli gaining a larger cortical representation (Rutkowski and Weinberger, 2005). In order to be certain that our optogenetic stimulation parameters did not induce plasticity that would confound interpretations of post-infusion data, we calculated the RMS values of sound-evoked CSD in AC. We examined the mean RMS of the CSD recorded from all channels on each shank, within 180 ms of sound onset where sound evoked significant CSD changes (Figure 5B). We analyzed the RMS values from the first 20 trials and the last 20 trials of the 100 trial pre-infusion session for all Chat-ChR2 transgenic mice and Ai32 control mice. If optogenetic stimulation induces plasticity effects, where the same sound stimulus will have a greater cortical representation, we would expect to see greater RMS values in sound-evoked CSDs. However, we found across both groups that there was no difference in the RMS values between control (n=6) and Chat-ChR2 animals (n=32) at the beginning (Wilcoxon rank sum test, p=0.707) or the end of the recording sessions (Wilcoxon rank sum test, p=0.701). These results suggest there were no plasticity effects on sound evoked auditory responses within our recording sessions as a result of optogenetic trials.

**Figure 5.**
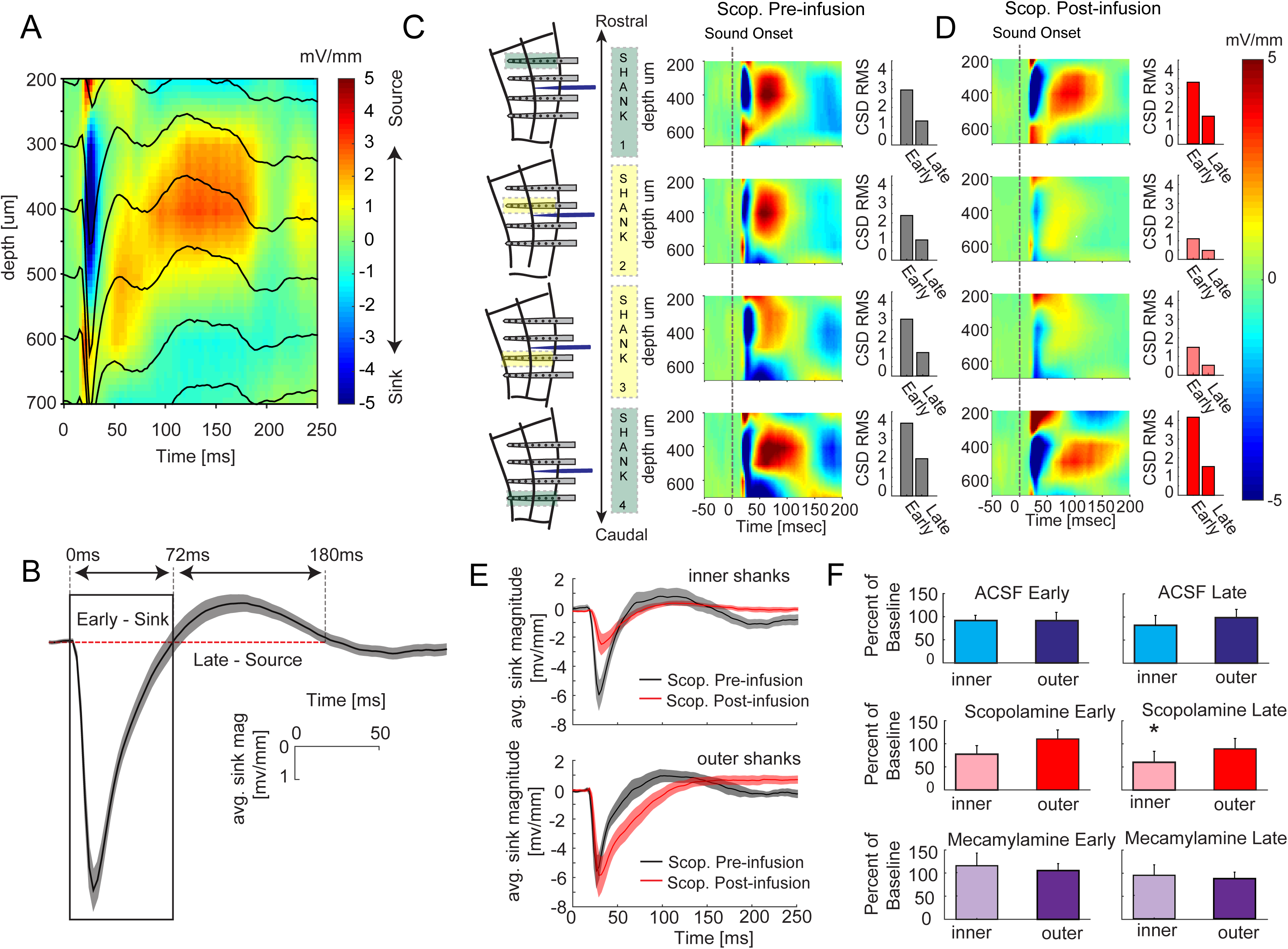
(Color): Effects of muscarinic blockade on CSD profiles in AC. **A:** ERPs in AC overlaid on CSD calculated from laminar ERP in a representative animal. **B:** Population sound-evoked CSD response from the channel exhibiting the largest thalamocortical response during pre-infusion period (mean ± s.e.m.). Red dotted line represents 0 mV/mm. Zero crossings were used to define an early window (0-72ms exhibiting granular layer current sinks) and a late window (72-186 ms exhibiting granular layer source). **C,D:** CSD responses across the four probes before (**C**) and after high-dose scopolamine infusion (**D**) separated by shank location. Bar graphs represent the calculated RMS across all channels of the CSD in the early and late windows as defined in **(B)**. Diagram illustrates recording site being shown and placement relative to infusion location in AC. Inner shanks (2&3) are closest to infusion pipette while 1&4 are furthest from the infusion site. Inner shanks are also highlighted yellow and outer shanks are highlighted green. Shank 1 is the most rostral placement while shank 4 is the most caudal. **E:** CSD response from the channel with the largest thalamocortical response from both inner and outer shanks before (black) scopolamine infusion and after (red) high dose scopolamine infusion (mean ± s.e.m). **F:** Change in net current of the CSD following cholinergic antagonist or ACSF infusions from the early (left) and late (right) windows. Bar height represents population mean, error bars are ± s.e.m. (*= p<0.05).

### 3.4 Muscarinic receptors contribute to intracortical signaling whereas nicotinic receptors alter the timing of thalamocortical input

To examine the effects of cholinergic receptor blockade on sound evoked AC responses across cortical layers, we characterized sound evoked CSD response profiles. In AC, CSD has been shown to allow for the dissociation of thalamic and cortical afferent pathways (Happel et al., 2010). The short latency phase of sensory-evoked CSD responses in AC consists of a current sink in the granular layers flanked by current sources in the infragranular and supragranular layers. This profile has been attributed to strong feedforward activation from the thalamus (thalamocortical), although within-column (intra-columnar) recurrent excitation in AC contributes to the overall amplitude of the early phase sink/source (Happel et al., 2010). Furthermore, the latent phase of the response consists of an inversion of the initial source-sink-source triplet, with a strong source in the granular layers and sinks in the supragranular and infragranular layers, primarily reflecting connections from adjacent regions of AC (intracortical). Similarly, in our dataset, we found that the auditory stimulus produced a large current sink in the middle layers ∼300-500 um from the cortical surface (Figure 5A, blue color), followed by a strong source that emerged ∼75 ms after auditory stimulus onset (Figure 5A, red color).

We computed CSD profiles for each recording shank, and determined the granular layers by identifying the channel with the earliest CSD sink. We then computed CSD profiles for each recording shank and used the largest CSD sink to characterize the time course of the sink source transition. The use of eight site probes, where sites were separated by 100 µm, allowed us to sample the majority of cortical depth, although infragranular locations were more sparsely sampled (see Figure S9 for quantifications of all electrode placements relative to the granular location). The population CSD from this channel, across animals, exhibited an early current sink (0-72ms) when the current was below zero, followed by a late current source (72-180ms) when the current was above zero (Figure 5B). We further determined the threshold of the early sink to estimate the response properties of AC to sound presentation. Across animals, the latency for current sinks, defined as significantly deviated from baseline, was 16.70±0.25 ms from stimulus onset, and the current sink returned to baseline at 59.07±3.73 ms (mean± s.e.m., n=128 shanks, 32 mice). These response profiles are consistent with general findings reported in AC using CSD analysis previously (Happel et al., 2010; Intskirveli and Metherate, 2012; Metherate et al., 2012; Szymanski et al., 2009; Szymanski et al., 2011).

To characterize the influence of cholinergic antagonists, we combined CSDs recorded from the two inner shanks, closer to the infusion pipette, and the two outer shanks, distal to the infusion pipette (Figure 4A). After administering high dose scopolamine, the magnitude of the CSD sinks and sources responses showed a reduction on the inner shanks that was not present on the outer shanks (Figure 5E).

We further investigated the effect of high dose scopolamine across all cortical layers by quantifying net laminar current, calculated as the mean RMS of the CSD recorded from all channels on each shank, during the two temporal windows described above: the early window (0-72ms), and the late window (72-180ms after noise onset) (Figure 5B). We utilized a non-parametric Friedman test to quantify the change in net current between drug conditions across shanks (inner/outer), time windows (early:sink/late:source), and animals. Across animals, we found that high dose scopolamine had a significant effect on net current (chi-square=5.9, p=0.02). Scopolamine produced moderate reductions in overall net laminar current flow on the inner shanks during the early window (Wilcoxon signed-rank test, p=0.055, n=9 mice) and robust effects during the late window (Wilcoxon signed-rank test, p=0.02, n=9 mice; Figure 5D, 5F). On the outer shanks, high dose scopolamine did not alter the net laminar current (Wilcoxon signed-rank test, p=0.91 for early, and, p=0.13 for late, n=9 mice; Figure 5D, 5F).

To further explore whether the effects of scopolamine on reducing current flow were specific to a particular layer or region, we analyzed the CSD profiles across the cortical laminae. We calculated the peak-trough magnitude of the CSD response for the supragranular, granular, and infragranular channels, and found that high dose scopolamine significantly reduced the peak-to-trough CSD magnitude in the supragranular and granular layers near the infusion site (supragranular, Wilcoxon signed-rank test, p=0.02; granular, Wilcoxon signed-rank test, p=0.04). All other layers and drug condition combinations failed to alter peak-to-trough CSD magnitude. This finding suggests that mAChR inactivation not only can reduce thalamocortical signaling in the granular layer, but also cortico-cortical signaling in supergranular layers. In comparison, neither ACSF, nor the nAChR blocker mecamylamine, changed net laminar current on the inner or outer shanks during the two windows analyzed (Wilcoxon signed-rank test, p> 0.05).

Finally, we examined the effects of the different cholinergic receptor antagonists on thalamocortical activation by analyzing the evoked CSD sink onset latency. Latency to sink onset (<20ms) is thought to be a more direct measure of thalamocortical activation as sink amplitude involves both contributions from thalamic input and intra-columnar recurrent excitation (Happel et al., 2010, Szymanski et al., 2011). Mecamylamine significantly delayed evoked response latency on the inner shanks, but not on the outer shanks (Wilcoxon signed-rank test, p=0.004, n=5 mice; Figure S2A-S2B, S2D), confirming that the dosage of mecamylamine was efficacious in blocking nAChRs, and consistent with delays reported previously at this concentration (Liang et al., 2006; Metherate, 2004). Given the strong influences of scopolamine on intra-columnar activity which contributes to sink amplitude described above (Figure 5E), we could not perform a statistical quantification of sink onset latency changes for high dose scopolamine. Because of this, we cannot rule out the possibility of contributions of mAChR to thalamocortical latency, although we found no differences in onset latency for low dose scopolamine or vehicle conditions on the inner or outer shanks (Wilcoxon signed-rank test, p> 0.05). Together, these findings demonstrate that timing of the initial sound-evoked response depends on intrinsic nAChR activation in our paradigm.

### 3.5 Scopolamine enhances the duration of sound evoked responses in AC outside of the infusion area

Interestingly, while high dose scopolamine infusion failed to influence net current on the outer shanks, it did moderately delay the sink-source transition by 28ms (Wilcoxon signed-rank test, p=0.07, n=9 mice; Figure 5D, 5E). To examine this more closely, we analyzed the time-course of the evoked MUAs in AC. Granular layer MUs showed the strongest onset responses, which returned to baseline within ∼ 75ms. Interestingly, high dose scopolamine did not alter the time course of MU responses in the granular layer on either the inner or outer shanks (Figure 6A-6B); however, MUs in the extragranular (supragranular and infragranular) layers were significantly broadened on the outer shanks and trended toward significantly broadening on the inner shanks (high-dose:inner, t(122)=-1.97; p=0.051, outer, t(121)=-4.07;p= 8.41 e-5; Figure 6C-6D). The broadened responses were also accompanied by an increase in the cross correlation of MUA between and within shanks at lags of 0-500ms (Figure S3A). Similarly, low-dose scopolamine also significantly broadened the spiking response in extragranular layers on the inner shanks (t(94)=-4.33; p=3.71e-5), and outer shanks(t(93)=-2.04, p=0.04), while having no significant effect in the granular layer (p>0.05). We saw no effects of vehicle or mecamylamine on MUs in any cortical layers (p>0.05; Figure 6B, 6D). We further analyzed the MUs from individual channels across all layers of AC to determine if any of the channels lost responsivity following high dose scopolamine infusion. While the population peak firing rate was maintained at sound onset when all channels were considered (Figure 6A-6C), a greater number MUs recorded from channels nearest the infusion cannula showed a reduction in evoked firing rate compared to the baseline (see methods; Figure S8A). In addition, following scopolamine infusion, approximately half of the MUs recorded in the supragranular layers on the outer shanks also showed a decrease in evoked responses, whereas MUs recorded in the granular or infragranular layers showed little change from the baseline condition.

**Figure 6.**
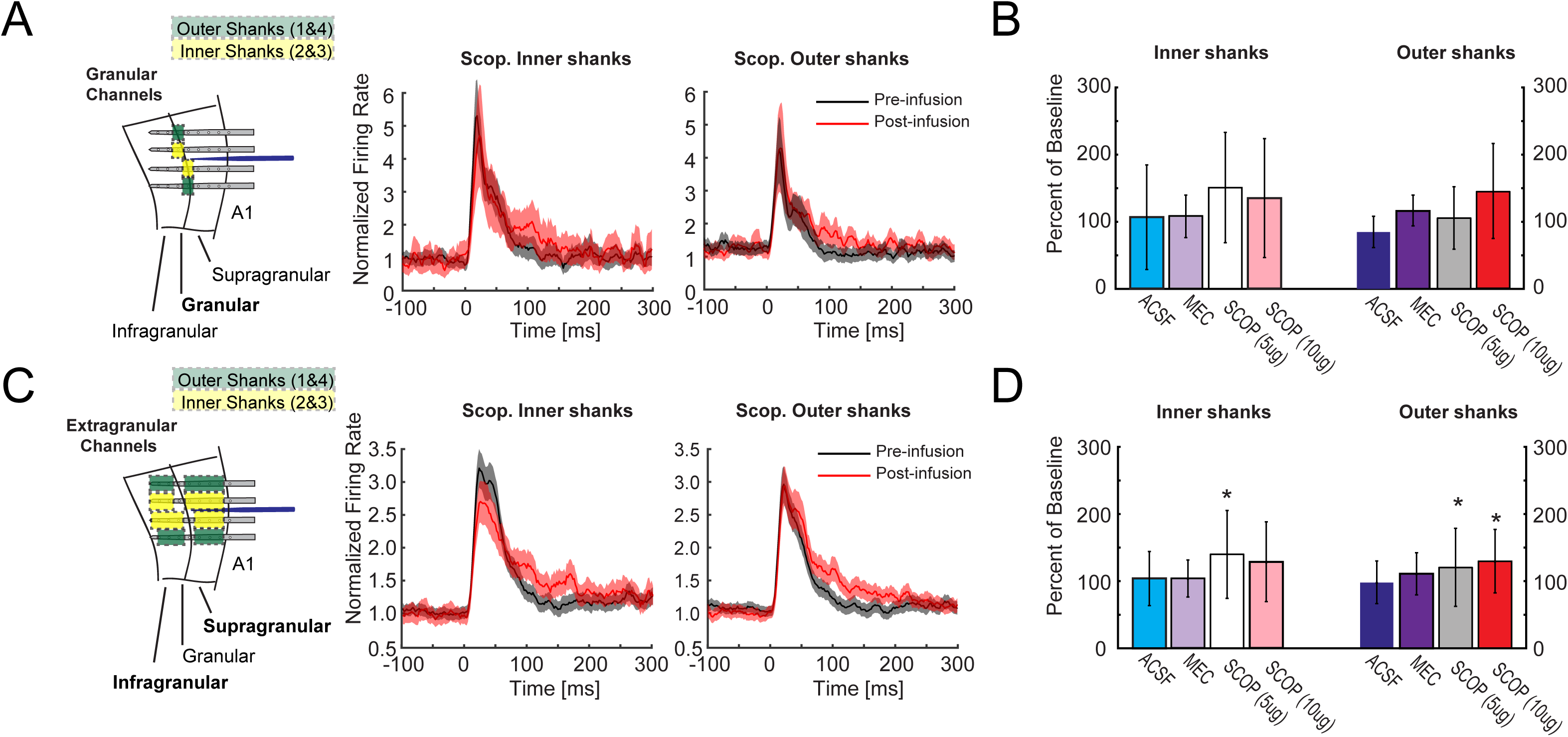
(Color): Endogenous muscarinic signaling modulates the strength of MUA in AC during sound presentation. **A:** Diagram illustrating recording/infusion configuration and location of signals shown on the right. Inner shanks are highlighted yellow and outer shanks are highlighted green. Granular channels are highlighted. Granular layer MU population responses from inner and outer shanks before (black) and after (red) high dose scopolamine infusion (right). **B:** Integrated MUA from the granular layers from the firing rate peak until 200ms into sound presentation (0-200ms). Bar graphs are changes normalized to the pre infusion baseline (* = p<0.05). **C:** Diagram illustrating recording/infusion configuration and location of signals shown on the right. Extragranular channels are highlighted. MUA from extragranular channels on the inner (left) and outer (right) shanks before (black) or after (red) high dose scopolamine infusion. **D:** Same as B, for extragranular layers. Figures are shown as population mean ± s.e.m. (* = p<0.05).

Together, these findings demonstrate that AC neurons exhibited reduced responsivity combined with prolonged current source activation at outer supragranular sites following blockade of endogenous mAChRs. This suggests that mAChR signaling during passive sound processing could contribute to the timing of intracortical auditory response dynamics in AC. It is interesting to note that the broadening of MUA is well timed to the delayed sink-source transition described earlier, providing further evidence of the effects of acetylcholine in sharpening auditory responses in AC through mAChRs.

### 3.6 Endogenous muscarinic activation in AC promotes evoked auditory responses in PFC

To determine how cholinergic receptors in AC influence sound evoked PFC responses, we characterized the effects of different antagonists on sound evoked ERPs in PFC. We found that a small local injection of high dose scopolamine into AC was sufficient to significantly reduced the strength of the mean ERP magnitude across all PFC recording sites (i.e., CG to MO subregions; Wilcoxon signed-rank test, p= 0.008, n=9 mice; Figure 7Aii, 7B). We observed no correlations between ERP amplitude reduction and recording depth (R=0.07, p=0.44, n=144 recording sites, 9 mice), suggesting that scopolamine broadly influences sound responses across all PFC subregions measured. Low dose scopolamine infusion produced a moderate, but non-significant decrease in PFC ERP magnitude (Wilcoxon signed-rank test, p= 0.22, n=7 mice, Figure 7B). Neither ACSF nor mecamylamine had any effect on PFC ERP responses (Wilcoxon signed-rank test, p>0.05, Figure 7Ai, 7B, S4A).

**Figure 7.**
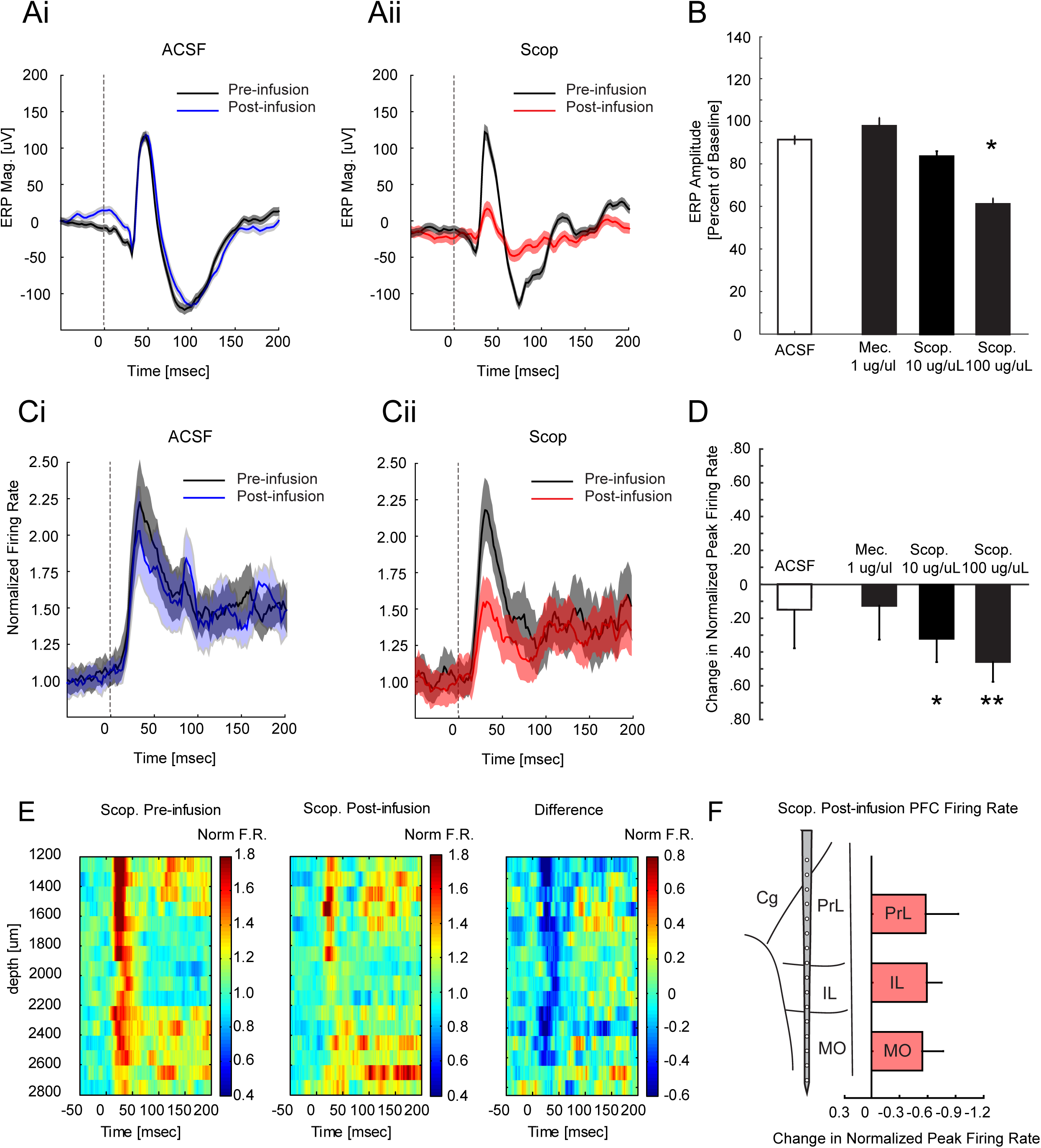
(Color): Scopolamine infusion in AC reduced ERP amplitude and spiking activity in PFC. **A:** Example sound evoked PFC ERP responses before (black) and after (blue) local infusion of ACSF (i) or high dose scopolamine (red) in AC (ii). Solid lines indicate mean, and shaded areas are mean ± s.e.m. **B:** ERP amplitude after infusion of different drugs normalized to pre-infusion amplitude. Error bars are ± s.e.m. **C:** Normalized MUA responses before (black) and after (red) infusion of ACSF (i) or high dose scopolamine (Cii). Lines are mean, and shaded areas are mean ± s.e.m. **D:** Comparison of the change in normalized MUA for all drug conditions. Bar graphs show mean ± s.e.m. (* = p<0.05; ** = p<0.01). **E:** Mean population PFC MU response across channels from the pre-infusion (left), post-Infusion (middle) and pre-post difference (right). F: Reduction in normalized firing rate by PFC subregion. All regions showed significant reductions from baseline that did not differ from one another. (Cg; Cingulate Cortex, Prl; Prelimbic cortex, IL; Infralimbic Cortex, MO; Medial Orbital Cortex).

Next, we examined the effects of drug infusion on PFC MU activity. Both high-dose and low-dose scopolamine infusion in AC robustly attenuated the sound evoked MU responses in PFC, with highdose scopolamine producing a reduction twice that seen with low-dose scopolamine (Figure 7C-7D, t(111)=2.26; p=0.026, high dose: t(137)=3.709; p=3.02e-4). Furthermore, this reduction was present across all PFC recording sites with no significant differences between PFC subregions (Chi-square=0.144, p=0.487, Figure 7E, 7F). Neither ACSF nor mecamylamine infusion had any influence on PFC MU activity (p>0.05, Figure 7D). Together, these results demonstrate that auditory responses in PFC are dependent upon activation of AC, and that endogenous muscarinic activation in AC is conducive to subsequent activation of PFC.

### 3.7 The role of cortical desynchrony on PFC auditory responses

One general concern we had was that animals might be expressing various levels of engagement throughout the passive presentation of sound, thereby influencing the level to which prefrontal networks are recruited. To address this, we characterized the strength of cortical desynchrony during the ITI as a proxy for engagement to the stimulus. Cortical desynchrony is associated with increased locomotion and generalized activity (Ma et al., 2017; Poulet and Petersen, 2008) and has been used as a measure of general arousal (Ma et al., 2017; Shinba et al., 2000). Given that cortical desynchrony has also been implicated in improving sensory responses (Goard and Dan, 2009; Kalmbach and Waters, 2014; Metherate and Ashe, 1993; Pinto et al., 2013), we assessed the strength of cortical desynchrony across mice, and across trials, to determine how the strength of cortical desynchrony preceding sound onset may influence the strength of the PFC response. We calculated the cortical desynchrony value, in the 750ms prior to sound presentation, and then divided the responses into two groups – those with cortical desynchrony values above the median, and those below the median (Figure S6B). When we analyzed the magnitude of the PFC ERP across the two conditions, we found no significant difference in the response between the two conditions (Supplemental Figure 6C; Wilcoxon rank-sum test p=0.879). While cortical desynchrony values varied slightly across trials, it did not have an impact on the strength of the PFC evoked response in mice that were passively presented with auditory stimuli. In addition, cortical desynchrony was significantly higher during optogenetic stimulation when compared to the desynchrony values measured during the ITI (Supplemental Figure 6A; (passive vs. stimulation, Wilcoxon sign-rank test, p=6.036e-6), suggesting desynchrony levels can be elevated from the baseline period via cholinergic stimulation

Taken together, this suggests that cortical desynchrony levels remained relatively constant across subjects within this study and were at levels lower than what we found in the same mice during optogenetic stimulation.

## 4. Discussion

This study aimed to determine if auditory signals are conveyed to multiple PFC regions during passive sound processing and whether cholinergic receptors in AC contribute to modulating auditory representations under these conditions. We performed simultaneous recordings in both structures while awake head-fixed mice were exposed to repeated delivery a white-noise stimulus in the presence and absence of cholinergic antagonists applied at concentrations shown to be effective in blocking cholinergic activation.

We found that PFC responds robustly to sensory input during passive presentation, exhibiting strong ERPs, and activating a significant proportion of the neurons across all medial PFC subregions recorded. We further found that responses in PFC were delayed by ∼10 ms from the onset of activation in AC, demonstrating that auditory information is represented in AC before being relayed to PFC through cortical or subcortical connections. These findings are similar to a recent study showing that another frontal region, orbitofrontal cortex, is also responsive to auditory input under passive presentation (Winkowski et al., 2017). The number of PFC units (∼12%) responsive to the white noise auditory stimuli in the passive condition in PFC is significantly less than what has been reported for prefrontal regions in aversive tasks such as fear learning where (25-75%) prefrontal neurons showed high sensitivity for the auditory conditioned stimulus following fear conditioning (Baeg et al., 2001; Burgos-Robles et al., 2009). Frontal cortex neurons can also be gated by task performance, becoming more sensitive to the same auditory stimuli when animals are introduced to the training context (Fritz et al., 2010). Our findings add to these bodies of work, by finding that a subset of prefrontal neurons track auditory stimuli during passive presentation in addition to being recruited, or gated, by cognitive tasks. The number of responsive neurons were also strikingly similar to the number of cells shown to be responsive for non-specific auditory activity in non-human primates where approximately 11% of single units showed broadband sensitivity to auditory input (Romanski et al., 2005).

We do not know the pathway by which auditory information is conveyed to PFC in our paradigm, however, connections between auditory association areas and prefrontal regions have been shown in both primates (Barbas, 2007; Romanski et al., 1999; Romanski and Goldman-Rakic, 2002) and rodents (Budinger et al., 2008; Martin-Cortecero and Nunez, 2016). We cannot rule out the possibility that auditory input is conveyed through subcortical structures (Hoover and Vertes, 2007), although the strong effect of local scopolamine infusion into AC reduces ERP responses in PFC by ∼70% suggests a primary auditory origin. This is consistent, with a recent study in anesthetized animals, where small local infusion of lidocaine into A1 effectively blocked PFC responses to auditory input (Martin-Cortecero and Nunez, 2016). In addition, that study found a delay of 8-18ms between AC and PFC, and is consistent with our findings of ∼10.5 seconds in this study and suggests the pathways active during anesthesia are likely the same pathways carrying sound information in awake animals under passive presentation. While it is possible that a 70dB SPL stimulus could be aversive and sound information could be conveyed via limbic or fear circuits, in fear conditioning studies where latency was examined, the robust increase in responsivity of prefrontal neurons for the auditory stimulus occurs ∼100ms after sound onset (Burgos-Robles et al., 2009). Presumably, if auditory information was conveyed via this circuitry, it might be expected to have a delayed presence in PFC relative to our findings.

On a related note, although we cannot directly assess the behavioral state of the animal in these studies, cortical desynchrony was robustly elevated by optogenetic stimulation of the NB, suggesting higher states of cortical desynchrony were achievable from the baseline condition. Interestingly, despite evidence that states of high cortical desynchrony can enhance processing in sensory cortices (Goard and Dan, 2009; Kalmbach and Waters, 2014; Metherate and Ashe, 1993; Pinto et al., 2013), we found no significant effect of cortical desynchrony levels on the magnitude of the PFC sound evoked response in our animals.

With regards to information processing in AC, we found significant effects of cholinergic antagonism on local processing. Since auditory stimuli are first relayed to AC and sound processing depends on interactions between thalamocortical inputs and cortico-cortical connections that are reflected in the laminar response profile (Happel et al., 2010), we also examined the effects of cholinergic antagonisms on CSDs in AC. Our study revealed that nAChR antagonism solely influenced the timing of the onset response to thalamic input into AC, delaying the sound evoked CSD response on the recording sites nearest the drug infusion. This demonstrates that we delivered a regionally localized effective dose and it produced an effect consistent with other local infusion mecamylamine studies in AC (Liang et al., 2006). Our finding that mecamylamine alters response latency provides evidence for an ongoing role of nicotinic signaling in sensory processing under basal conditions consistent with previous findings (Metherate, 2004). Based on previous studies (Intskirveli and Metherate, 2012; Kawai et al., 2007), we expected that mecamylamine may attenuate the auditory evoked ERP at short time scales but we found no evidence of this during passive processing of sound. It is likely that the influence of nicotinic activation on sound processing is altered by attentional demand and the strength of cholinergic tone. For example, acetylcholine release can improve sensory encoding by enhancing responses to relevant stimuli during sensory discrimination (Liang et al., 2008). It is interesting that in the Liang et al. study, the nAChR agonist nicotine lowered response thresholds for characteristic frequency stimuli and increased ERP amplitude only in rats performing at high levels in a behavioral task. In intermediate and poorly performing animals, nicotine only influenced the onset latency. Our results in naïve non-performing animals were similar; the only observable effect of mecamylamine was to delay onset latency. These findings further provide evidence that basal levels of acetylcholine contribute to signal processing in passive sound processing, although the strength of regulation is most likely dynamic and heavily influenced by cognitive state.

In contrast, mAChR antagonism strongly influenced intracortical signaling within AC. The mAChR antagonist scopolamine had a strong attenuating effect on the magnitude of the late phase CSD response, and the amplitude of the early phase response, but not on the thalamocortical sink onset latency at doses where latency could be measured. This suggests that mAChRs play a prominent role in modulating AC responses that are modulated through cortical interactions. We found that the reduction in net laminar current was coincident with sustained periods of evoked MUA on shanks distal to infusion, as well as a delayed sink-source transition following sound onset. The mechanism by which mAChRs produce this effect is still unknown, but could involve contributions of enhanced excitability of pyramidal networks or reduced inhibition of interneurons. One possible explanation is that input originating from adjacent auditory frequency response areas that would be inhibited, is attenuated in the presence of scopolamine, consistent with a mechanism where reduced lateral inhibition alters the shape of the time-course of sensory responses in AC (Wehr and Zador, 2003; Wu et al., 2008). This is consistent with the more general idea that broadly tuned cortical interneurons enhance the contrast of the auditory representations through horizontal interactions (Li et al., 2014). In our study, it is possible that scopolamine alters efferent inhibitory signaling from the region of the infusion and that the remaining inhibitory current takes longer to balance out excitatory drive from the thalamus, leading to a longer duration current sink. While we did not test this mechanism in this study, cortical frequency tuning is thought to depend on lateral inhibition, which enhances the contrast of frequency representation in AC. This process is critically dependent on the activity of L2/3 PV and SOM interneurons (Kato et al., 2017; Li et al., 2014), which can be strongly regulated by acetylcholine (Chen et al., 2015). Lateral Inhibition has been shown to be modulated by attention in humans (Engell et al., 2016), suggesting cholinergic signaling could play a pivotal role in shaping sensory response profiles across the AC tonotopy.

Our finding that scopolamine infusion into AC strongly reduced responsivity in PFC also suggests that mAChR activation contributes to circuit dynamics that are permissive to cortico-cortical gating and the elevation of sensory information into prefrontal networks. One potential mechanism by which acetylcholine could influence the strength of information flow, is by effectively lowering the threshold for excitation and promoting feedforward excitation by inactivating m-currents in layer II/III low threshold spiking interneurons. These neurons have been proposed to gate intracortical signaling by inhibiting fast spiking interneurons (Chen et al., 2015; Lee et al., 2015). This interpretation is supported in part by data showing a longer delay to the sink-source transition and prolonged time-course of MU responses. It is possible this is a result of reductions in recurrent interactions at superficial layers of AC reducing the late phase source magnitude (Figure S8B). This reduction in superficial current magnitude combined with a reduction in MU responses specifically in these layers supports a model where auditory output is supported by superficial layer efferents that ultimately contribute to the strength of the signal that reaches the PFC.

Previous studies have demonstrated that during high attentive states, regions of the auditory cortical tonotopy interact intracortically to enhance attended, and suppress unattended auditory stimuli (O’Connell et al., 2014). Interestingly, our results reveal that mAChRs play a prominent role in intracortical signaling within AC and long range cortico-cortical signaling between AC and PFC during passive sound presentation, which is complementary to those functions previously reported during attention tasks. Our findings suggest acetylcholine signaling enhances sensory processing indiscriminately, allowing bottom-up inputs to be refined and relayed to higher cortical areas for additional processing. This finding is consistent with a model in which PFC monitors stimuli that may later become important (i.e., behaviorally relevant cues used to guide behavior). Taken together, our data provide direct evidence that cholinergic mechanisms play a role in ascending auditory processing and are complementary to the roles ascribed for acetylcholine in facilitating top-down mediated attentional performance (Guillem et al., 2011; Hasselmo and Sarter, 2011; Sarter et al., 2005).

This previously unidentified requirement for acetylcholine in regulating long-range cortical signaling during passive sound processing also expands canonical views of the role of prefrontal cortex in sensory processing and cholinergic function. Acetylcholine may promote long-range network communication by enhancing intrinsic oscillatory patterns during stimulus processing (Roopun et al., 2010). In this regard, cortical projecting cholinergic systems may contribute to signal processing by gating information flow between interconnected cortical regions, in addition to strengthening thalamo-cortical signaling. It is interesting that scopolamine blocked the reduction in low frequency power (1-10 Hz) during optogenetically evoked cortical desynchrony without altering high frequency power (10-100 Hz). Combined with the fact that scopolamine also reduced intracortical signaling, it is possible that a role of muscarinic activity is to reduce low frequency power during periods where intracortical signaling is prioritized.

It is not known how mAChRs contribute to intracortical signaling or low frequency desynchronization during cholinergic release. More work will be necessary to determine if these two phenomena might be related, but cholinergic signaling of interneuron sub-types potentially contribute to both processes. For example, parvalbumin positive interneurons have been shown to contribute to feedforward inhibition within cortical circuits that mediate lateral suppression during auditory stimulation (Li et al., 2014), and a recent study suggested that superficial somatostatin positive interneurons may play an active role in cholinergic modulation of cortical desynchrony and neuronal decorrelation associated by shifting intracortical interactions (Chen et al., 2015). Future experiments will be necessary to determine how this process is accomplished and how distinct cholinergic receptor activity contributes to signal processing and cortico-cortical communication differently depending on the demands on attention.

## Acknowledgements

We thank members of the Han Lab for suggestions on the manuscript. We would also like thank Spencer Torene, Erik Roberts, Grant Fiddyment, and Wei Guo for their useful insights on data analysis and collection. X.H. acknowledges funding from National Institutes of Health Director’s new innovator award (1DP2NS082126), NIH 1R01NS087950-01, Pew Foundation, Alfred P. Sloan Foundation, and Boston University Biomedical Engineering Department. The authors declare no competing financial interests. Correspondence should be addressed to Xue Han.

## Author Contributions

N.M.J. and H.J.G. performed all experiments. N.M.J. processed and analyzed the data. K.S. and N.K. edited the manuscript and contributed to the interpretation of the results. Additionally, K.S. and N.K. provided technical support. X.H. supervised the study. N.M.J., H.J.G., and X.H. wrote the manuscript.

## Supplemental Figures

**Supplemental Figure 1.**
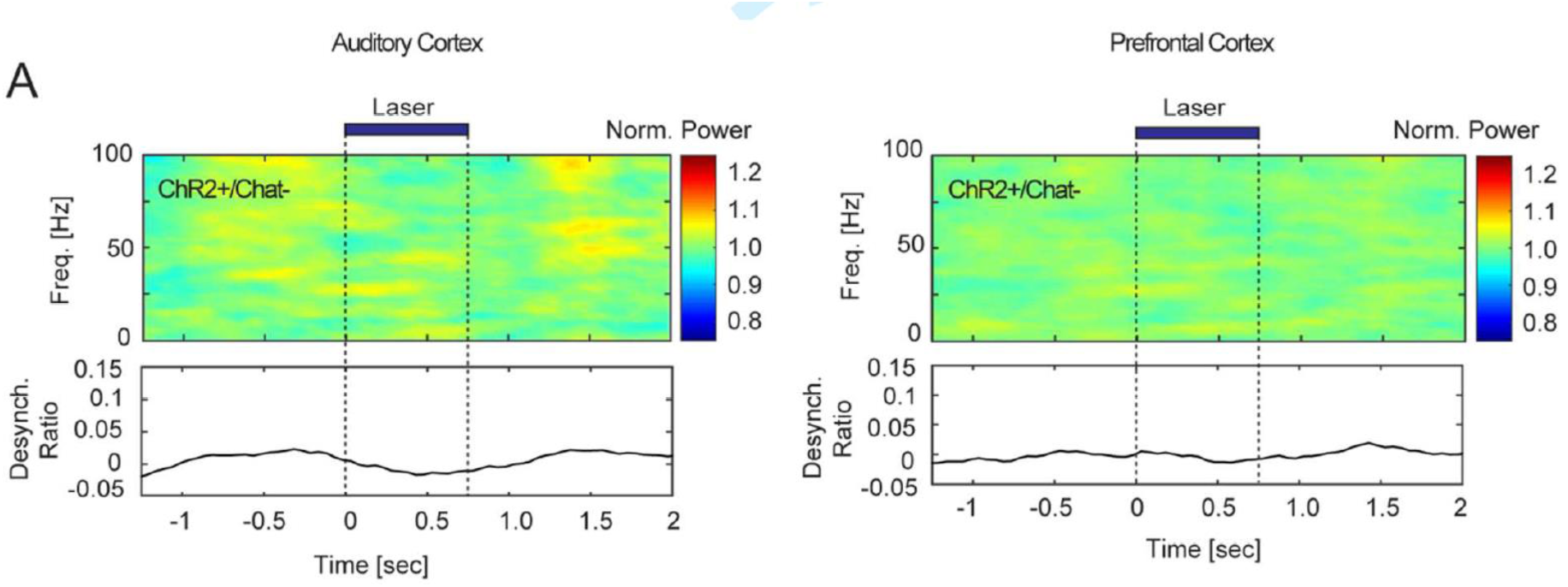
(Color): Laser light delivery to NB in Ai32 control mice (ChR2^+^/Chat^-^) does not produce cortical desynchrony. **A:** Trial averaged, baseline normalized spectrograms from AC (left) or PFC (right) during optical stimulation. The ratio of low frequency power (1-10 Hz) to high frequency power (10-100 Hz) for AC and PFC is shown below the spectrograms (bottom).

**Supplemental Figure 2.**
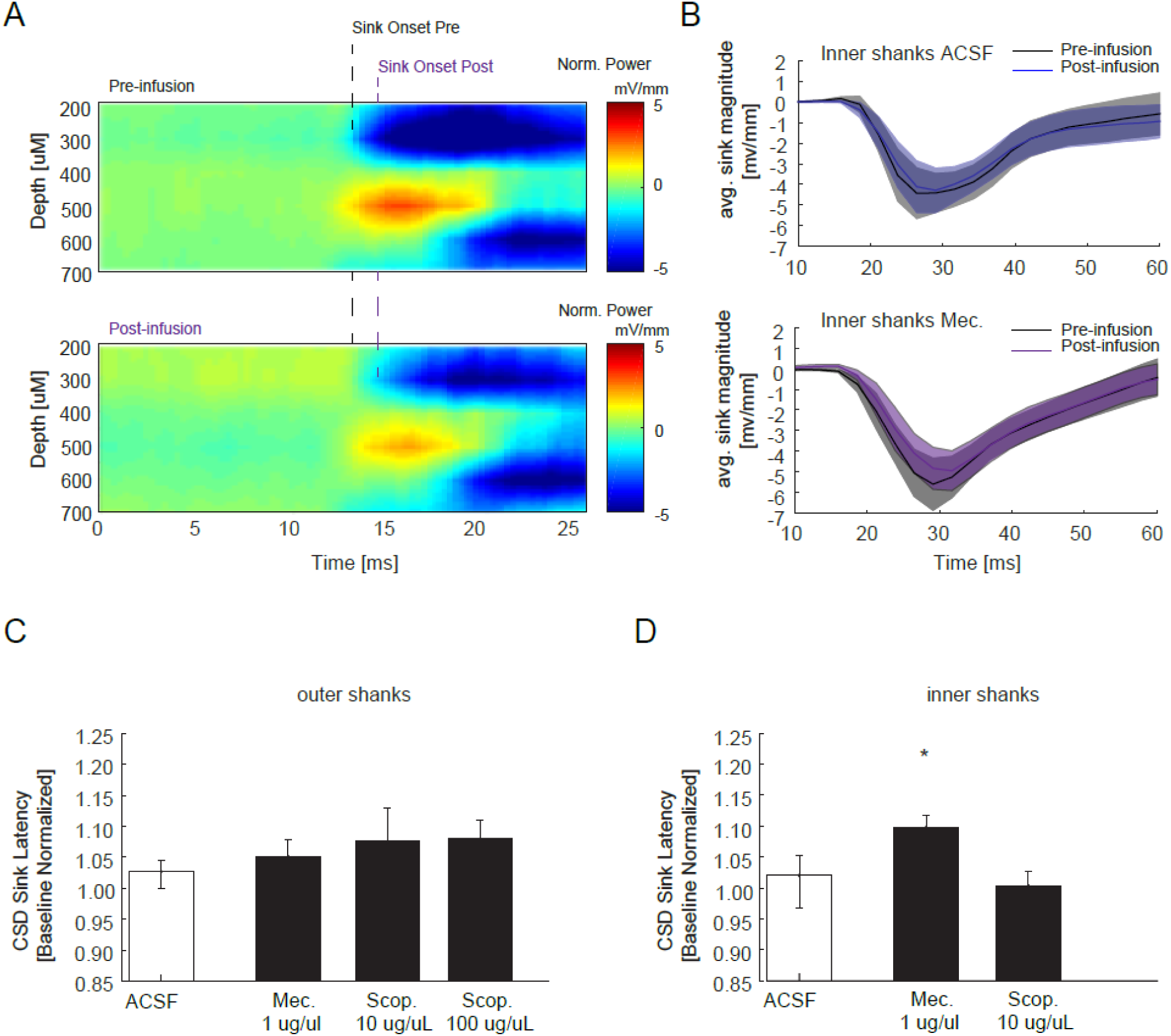
(Color): Mecamylamine delayed CSD current sink onset latency in AC. **A:** Representative CSD profile from an animal before (top) and after infusion of mecamylamine (bottom). Onset latency of thalamo-cortical sink is demarcated by the dashed line in the pre and post infusion periods. **B:** Population CSD before (black) and after infusion of vehicle (top, blue) or mecamylamine (bottom, purple) for the inner shanks (mean±s.e.m.). **C, D:** Population CSD latency for all drug conditions and vehicle normalized to baseline for outer shanks **(C)** and inner shanks **(D)** (mean ± s.e.m). Note for inner shanks the attenuation of the current sink amplitude prevented calculation of onset latency for the high dose scopolamine drug condition (* = p<0.05).

**Supplemental Figure 3.**
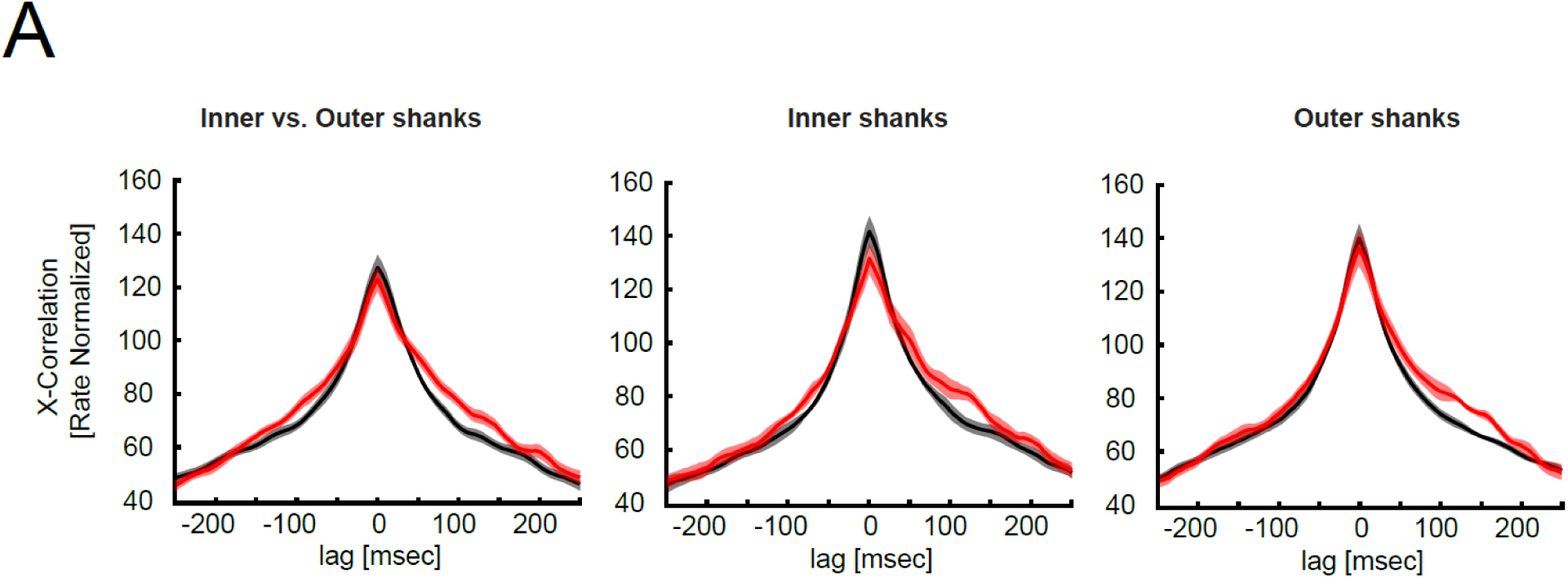
(Color): High-dose scopolamine increased the duration of cross correlation between evoked MUs in AC. **A: (left):** Cross correlation between shanks nearest scopolamine infusion site (inner shank) and those further away from infusion site (outer) from the sound period. **(middle and right):** Cross correlations computed between inner shanks (middle) and outer shanks sites (right) separately. Error bars are ± s.e.m.

**Supplemental Figure 4.**
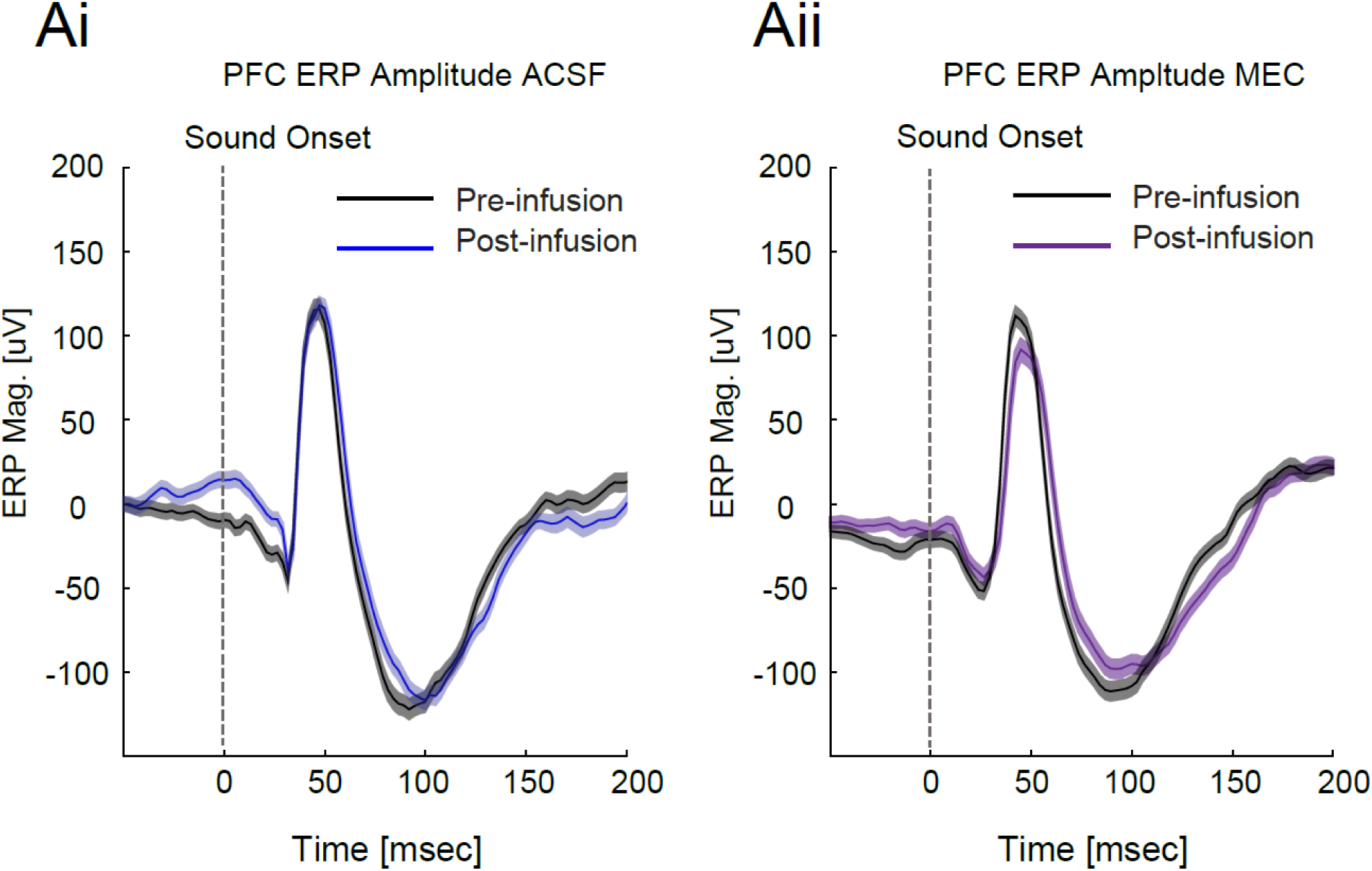
(Color): Mecamylamine infusion in the AC did not alter ERP amplitude in PFC. Example sound evoked PFC ERP responses before (black) and after local infusions of ACSF (Ai: blue) or mecamylamine (Aii:purple) into AC shown as mean ± s.e.m. Vertical dashed lines at zero denotes sound onset.

**Supplemental Figure 5.**
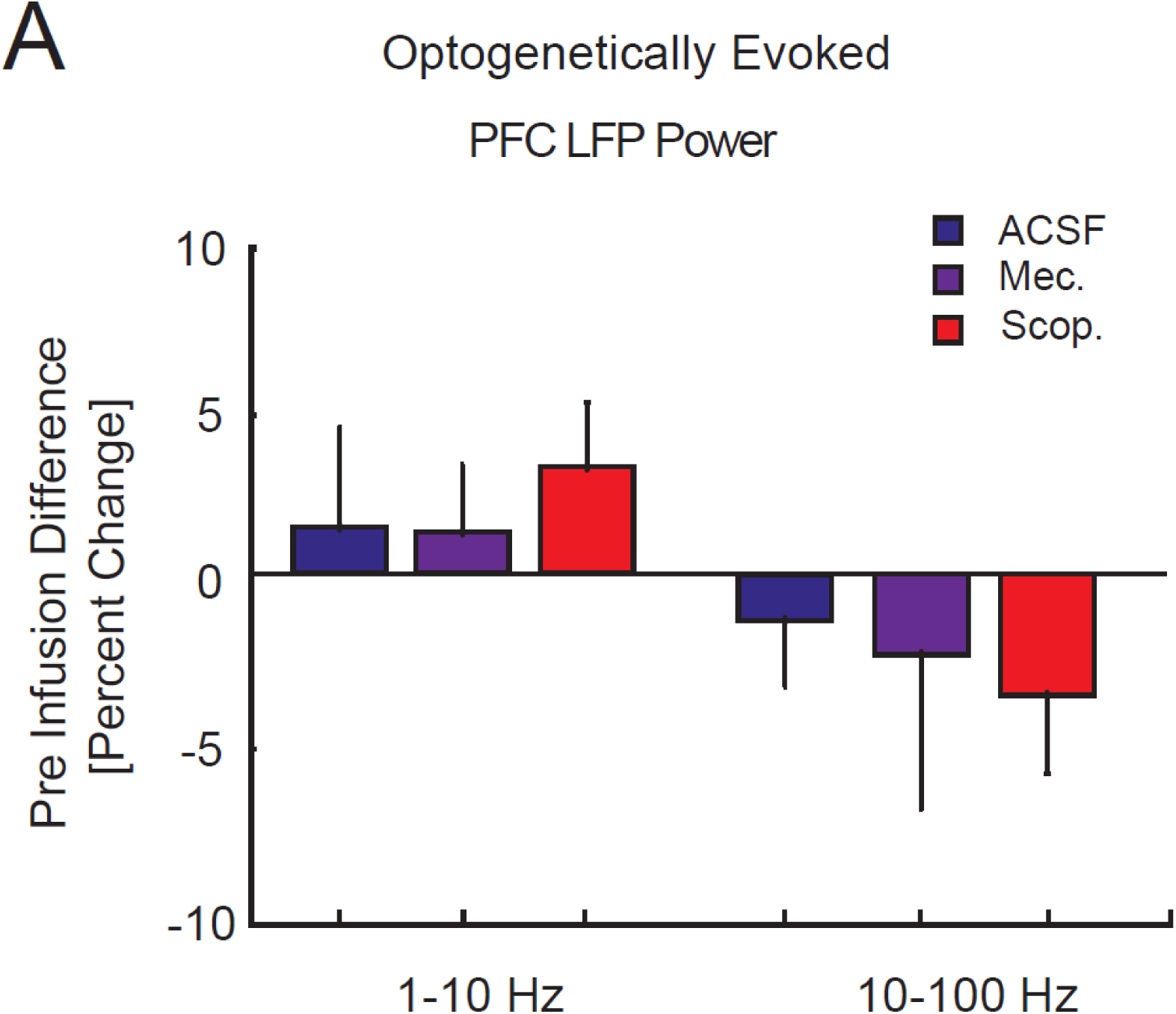
(Color): Scopolamine infusion in AC did not alter cortical desyncrony observed in PFC. **A.** Change in PFC LFP power by frequency range during optogenetic stimulation in the presence of antagonists or vehicle. Values are expressed as percentages of pre infusion power (mean ± s.e.m). Note that there were no changes in power (difference from 0) in either frequency band for PFC, despite large changes in power at the low frequency band for AC (see Figure 4E).

**Supplemental Figure 6.**
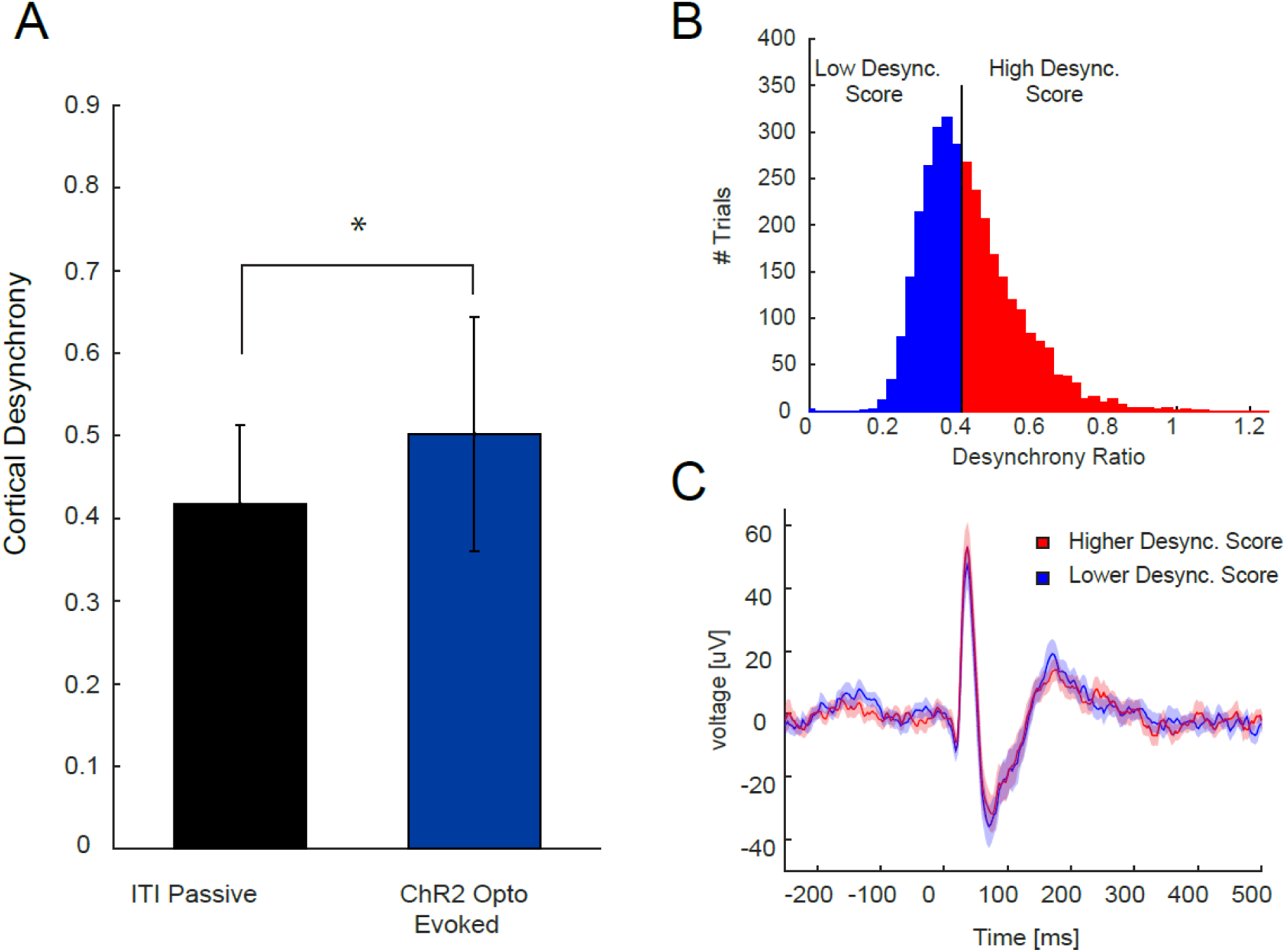
(Color): Evoked ERPs are independent of variations in cortical desynchrony level at the time the auditory stimulus occurs. **A.** Comparison of the cortical desynchrony values measured from the 750ms immediately before sound presentation, to the cortical desynchrony values during 750ms window of optogenetic stimulation of NB. Values are absolute ratio of low frequency power (1-10 Hz) to high frequency power (10-100 Hz). Data are reported as mean ± s.e.m (Wilcoxon sign-rank test, passive vs. stimulation, p=6.036e-6; ***= p<0.001). **B.** Binned histogram of cortical desynchrony values for all trials, across all animals, from the 4-9 second window of the ITI, preceding sound presentation. Median value is shown as a black line. Trials below the median were classified as having low cortical desynchrony levels (shaded blue), whereas trials above the median were classified as high cortical desynchrony levels (shaded red). **C.** Comparison of the PFC ERP on trials with low (color: blue) versus high (color: red) cortical desynchrony levels.

**Supplemental Figure 7.**
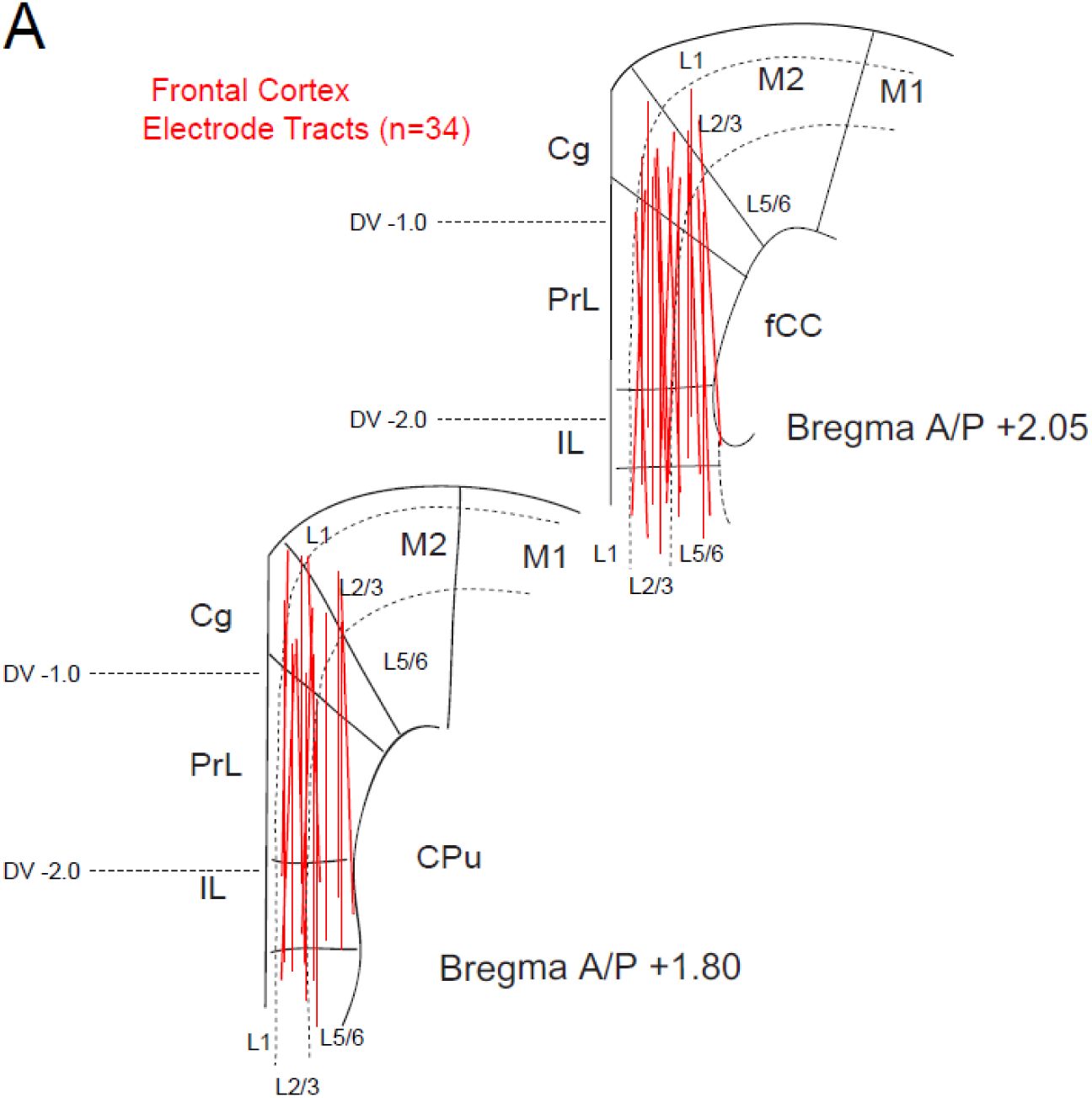
(Color): Histological reconstruction of electrode placement. Anatomical depiction of electrode placements of the 16-channel electrode array targeted to frontal cortex. Reconstructions are from 34 animals represented across two frontal (PFC) sections that best represent their anterior-posterior location. PFC penetration locations were ambiguous in four animals. Depth is noted as a D/V division as well as approximate layer boundaries between superficial and deep layers. **Abbreviations:** Cg; Cingulate Cortex, Prl; Prelimbic cortex, IL; Infralimbic Cortex, CPu; Caudate Putamen, M1; Primary Motor Cortex, M2; Secondary Motor Cortex.

**Supplemental Figure 8.**
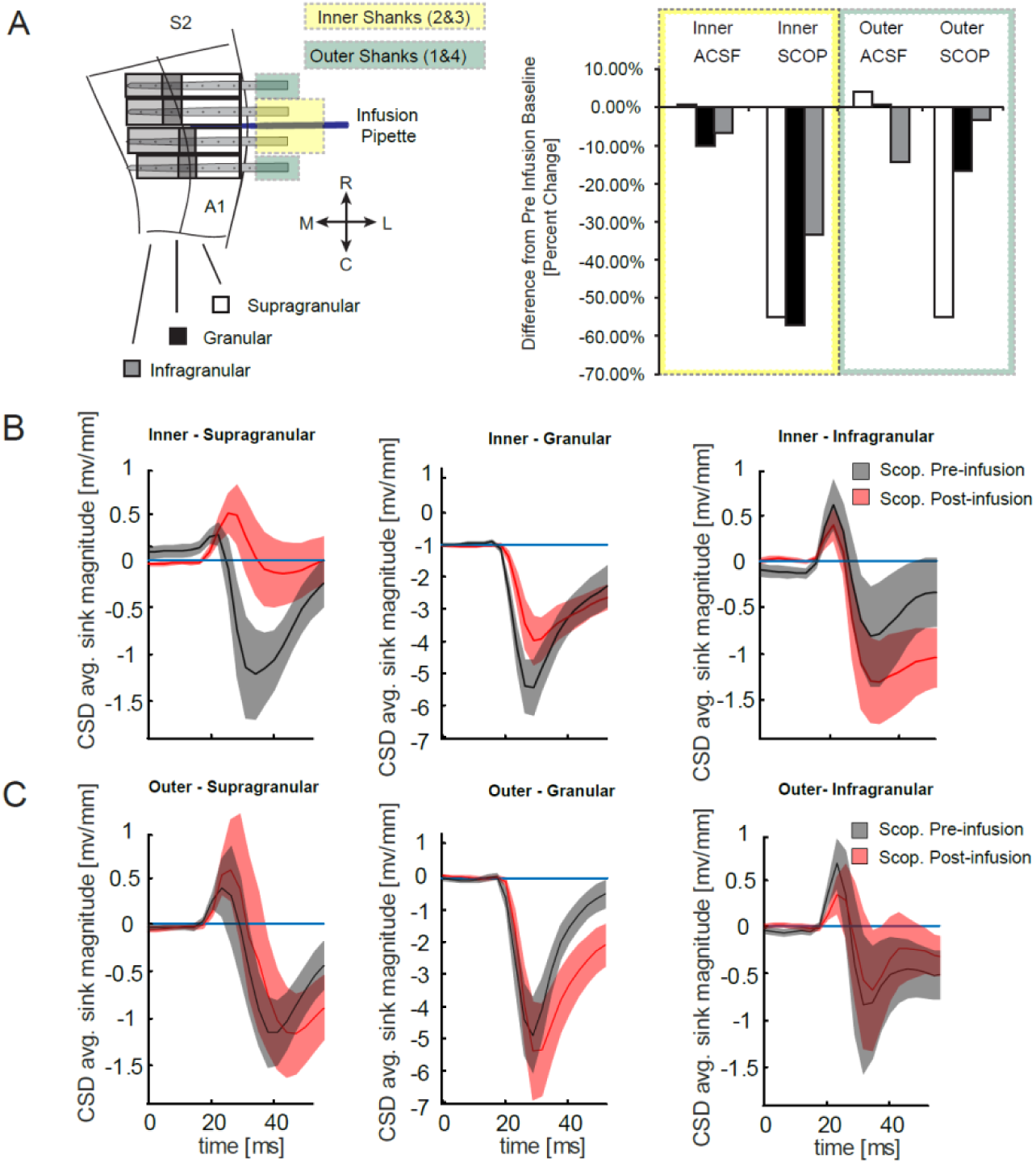
(Color): Neural responsivity and layer specific quantification of current flow. **A:** Neural responsivity following high-dose scopolamine infusion is separated by cortical layers. Diagram illustrating recording/infusion configuration in AC. Dotted boundary reflects dissociation between inner and outer shanks (left). Number of auditory responsive channels sorted by layer normalized to the number of channels responsive pre-infusion (granular: pre inner n=14/18, pre outer n=12/18, supragranular: pre inner n=20/80, pre outer n=20/65, infragranular: pre inner n= 30/43, pre outer n= 31/57). **B, C:** Population sound-evoked changes in CSD RMS sink minus source magnitude by shank location was significantly altered following scopolamine infusion (Chi-square=5.92, p=0.015). CSD pre-infusion responses are shown in black, and post-infusion responses are shown in red with inner shanks represented in **(B)** and outer shanks represented in **(C)**. Post-hoc analysis revealed significant attenuation for granular (Wilcoxon signed-rank test, p=0.04) and supragranular layers on the inner shanks (Wilcoxon signed-rank test, p=0.02).

**Supplemental Figure 9.**
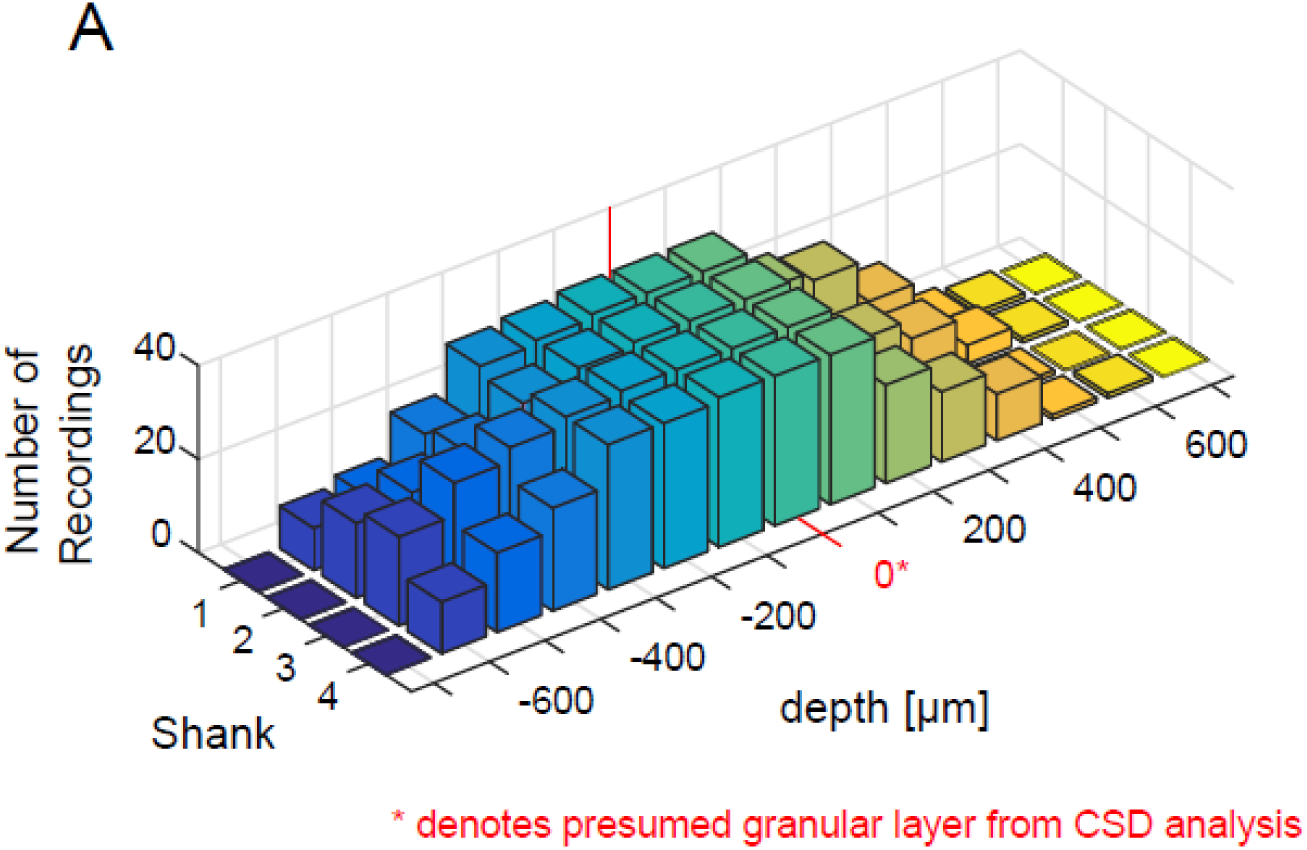
(Color): Distribution of recording sites separated by anatomical depth. The number of recording sites across all animals (n=34), plotted relative to the granular layer (shown as zero in red). Each shank is highlighted (x-axis) along with absolute depth (z-axis). Infragranular layers are shown as positive numbers, and supragranular layers are denoted with negative numbers. Note that the distribution of recording sites across shanks were consistent with supragranular layers having the largest proportion of represented channels.

**Supplemental Figure 10.**
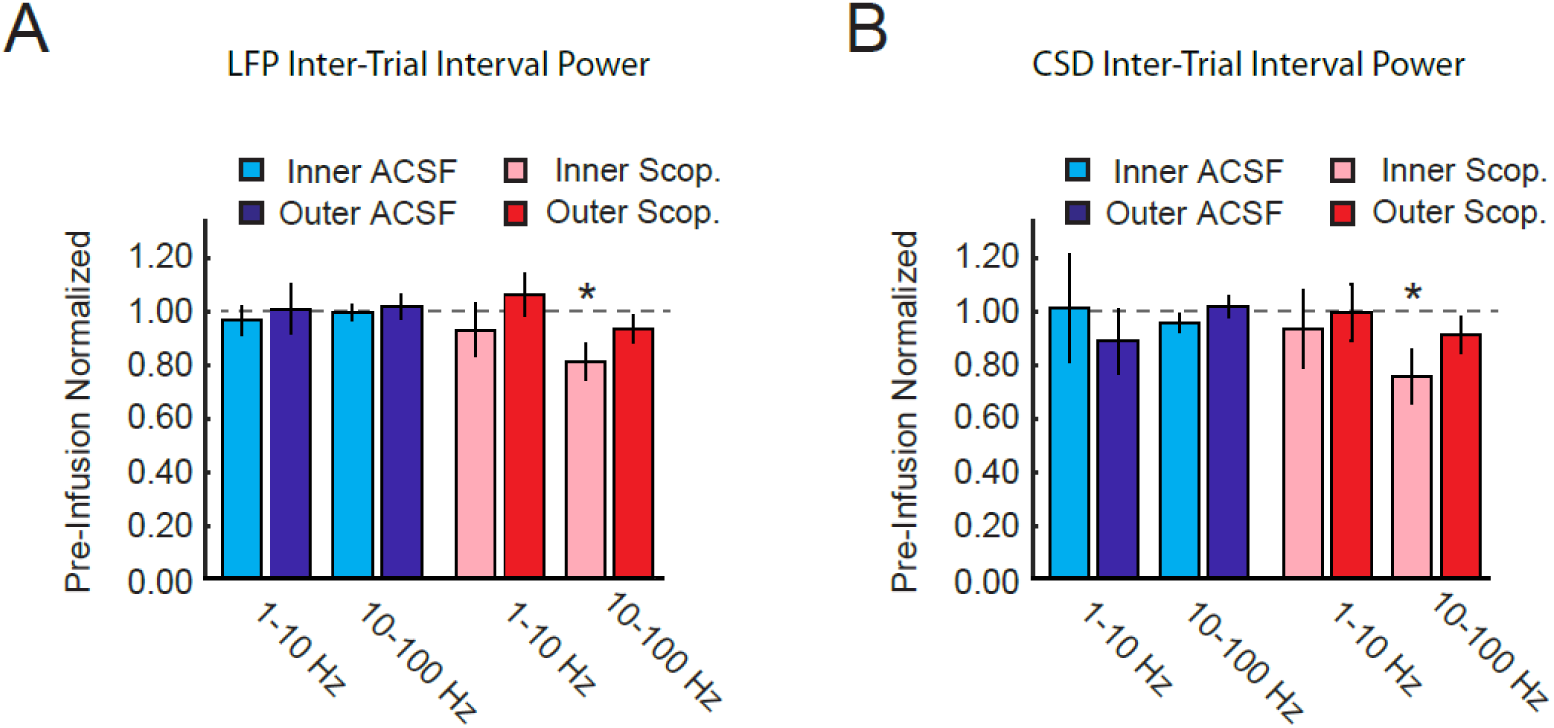
(Color): Comparison between LFP and CSD power at low or high frequencies during inter-trial interval. **A:** Population mean ± s.e.m normalized LFP power in the 1-10Hz frequency range and the 10-100Hz frequency range, on inner and outer shanks in AC, before and after infusion of ACSF (blue) or scopolamine (red). This same figure is also shown in figure 4B. **B:** Population mean ± s.e.m normalized CSD power in the 1-10Hz frequency range and the 10-100Hz frequency range on inner and outer shanks in AC before and after infusion of ACSF (blue) or scopolamine (red) for comparison to **(A)**. Note that only the 10-100Hz frequency range was reduced following scopolamine infusion in both analysis (*= p<0.05).

